# Lack of blocking activity in anti-CTLA-4 antibodies reduces toxicity, but not anti-tumor efficacy

**DOI:** 10.1101/2021.07.12.452090

**Authors:** Erica L. Stone, Kyle P. Carter, Ellen K. Wagner, Michael A. Asensio, Emily Benzie, Yao Y. Chiang, Garry L. Coles, Chelsea Edgar, Bishal K. Gautam, Ashley Gras, Jackson Leong, Renee Leong, Vishal A. Manickam, Rena A. Mizrahi, Ariel R. Niedecken, Jasmeen Saini, Savreet K. Sandhu, Jan Fredrick Simons, Kacy Stadtmiller, Brendan Tinsley, LaRee Tracy, Nicholas P. Wayham, Yoong Wearn Lim, Adam S. Adler, David S. Johnson

## Abstract

Anti-CTLA-4 antibodies such as ipilimumab were among the first immune-oncology agents to show significantly improved outcomes for patients. However, existing anti-CTLA-4 therapies fail to induce a response in a majority of patients and can induce severe, immune-related adverse events. It has been assumed that checkpoint inhibition, i.e., blocking the interaction between CTLA-4 and its ligands, is the primary mechanism of action for ipilimumab. In this study we present evidence that checkpoint inhibition is not a primary mechanism of action for efficacy of anti-CTLA-4 antibodies. Instead, the primary mechanism for efficacy is FcR-mediated Treg depletion in the tumor microenvironment. First, we identified a monoclonal antibody (mAb) that binds to CTLA-4 at an epitope that differs from ipilimumab’s by only a few amino acids, yet has limited checkpoint inhibitor activity. Surprisingly, the weak checkpoint inhibitor has superior anti-tumor activity compared to ipilimumab in a murine model. The weak checkpoint inhibitor also induces less Treg proliferation and has increased ability to induce *in vitro* FcR signaling and *in vivo* depletion of intratumoral Tregs. Further experiments showed that the enhanced FcR activity of the weak checkpoint inhibitor likely contributes to its enhanced anti-tumor activity. Importantly, we also showed that weak checkpoint inhibition was associated with lower toxicity in murine models. Our work suggests that new anti-CTLA-4 drugs should be optimized for Treg depletion rather than checkpoint inhibition.

## INTRODUCTION

The FDA-approved anti-Cytotoxic T-Lymphocyte-Associated protein 4 (CTLA-4) antibody ipilimumab induces long-term, durable responses in a subset of patients (1, 2), and is a mainstay of immune-oncology (IO) therapy. Ipilimumab combined with nivolumab (an anti-PD-1 antibody) is an FDA-approved first-line therapy for advanced melanoma and intermediate or high-risk renal cell carcinoma (3, 4). Additionally, ipilimumab plus nivolumab is effective in patients with cancers that are typically resistant to anti-PD-1 monotherapy, including melanoma patients with brain metastases (5, 6). However, ipilimumab has a relatively low response rate (10-16%) (1, 2, 7, 8, 9) and can induce severe adverse events including colitis, dermatitis, hepatitis, endocrinopathies, and renal dysfunction (1, 2, 3, 4). Thus, patients would benefit from an improved anti-CTLA-4 therapy with enhanced efficacy and/or reduced toxicity.

The mechanism of action of anti-CTLA-4 drugs remains controversial (10). CTLA-4 monoclonal antibodies (mAbs) such as ipilimumab were originally intended to block the binding of CTLA-4 to its ligands, the B7 proteins CD80 and CD86, i.e., “checkpoint inhibition”. Blocking CTLA-4 binding to B7 proteins frees B7 proteins to bind to CD28, inducing T cell co-stimulation and activation (11). However, acute deletion of *Ctla4* does not limit growth of MC38 tumors in a murine model (12), suggesting that loss of CTLA-4 activity alone may not be sufficient to enhance anti-tumor immunity. In addition to blocking the interaction of CTLA-4 with its B7 ligands, anti-CTLA-4 mAbs are also able to induce antibody-dependent cell-mediated cytotoxicity (ADCC) and antibody-dependent cellular phagocytosis (ADCP) of intratumoral FOXP3^+^ regulatory T cells (Tregs) (13, 14, 15, 16, 17, 18, 19), which express comparatively high levels of surface CTLA-4 (14, 15, 16, 17). It is reasonable to believe that these activities contribute to the anti-tumor activity of CTLA-4 mAbs, because it is known that depletion of intratumoral Tregs induces anti-tumor immune activity (20). Furthermore, several studies in murine models showed that Fc effector function and/or activating Fc receptors are necessary for the anti-tumor activity of anti-CTLA-4 (14, 15, 16, 17, 18). Importantly, intratumoral Treg depletion correlates with efficacy of ipilimumab in patients with melanoma (13, 14).

Despite their ability to deplete intratumoral Tregs, conventional CTLA-4 antibodies have been shown to enhance peripheral Treg levels or proliferation in murine models, cynomolgus monkeys, and patients (16, 17, 21, 22, 23). The difference in the effect of anti-CTLA-4s on peripheral and intratumoral Tregs is likely due to the amount of CTLA-4 expressed on the surface of Tregs in the periphery versus the tumor. In murine models and patients, intratumoral Tregs express more surface CTLA-4 (15). As the ability of antibodies to mediate depletion correlates with the amount of antibody on the cell surface (24), and even intratumoral Tregs have relatively low surface CTLA-4 levels (14, 15, 16), it is likely that only intratumoral Tregs express enough surface CTLA-4 for anti-CTLA-4 mAbs to mediate depletion via ADCC/ADCP. While CTLA-4^Lo^ cells, including effector T cells and peripheral Tregs, do not express enough surface CTLA-4 to be depleted in the presence of anti-CTLA-4, their surface CTLA-4 expression is sufficient to allow anti-CTLA-4s to enhance their co-stimulation via checkpoint inhibition. Further evidence that CTLA-4 acts to limit Treg proliferation comes from experiments in which *Ctla4* was acutely deleted in adult mice, which resulted in enhanced Treg proliferation and accumulation (12). One explanation for this observation could be that co-stimulation is not only important for activation of effector T cells, but it also enhances the activation and activity of regulatory T cells (25, 26). Thus, just like first generation IL-2 therapy (27, 28, 29), first generation anti-CTLA-4 therapies may preferentially induce Treg proliferation, thus limiting the efficacy of these therapies.

The contribution of various mechanisms of action of anti-CTLA-4 therapies to the severe adverse events induced by these therapies is also not well understood. However, it is known that *CTLA4* haploinsufficiency leads to inflammatory symptoms reminiscent of the adverse events induced by ipilimumab and tremelimumab (30, 31, 32, 33, 34, 35, 36). In agreement with this, acute deletion of *Ctla4* in adult mice can lead to lymphoproliferation and immune pathology (37). Furthermore, it has recently been shown that immune checkpoint blockade induction of colitis in melanoma patients is *not* associated with a loss of Treg cells (38). Taken together these data suggest that blocking the interaction of CTLA-4 with its B7 ligands may have a considerable contribution to the adverse events induced by conventional anti-CTLA-4 therapies.

Thus, a possibility is that an anti-CTLA-4 that induces intratumoral Treg depletion but has little ability to block the interaction of CTLA-4 with CD80/CD86 would have improved anti-tumor activity while limiting the development of adverse events. Here we show that compared to ipilimumab, the weak B7 ligand blocking anti-CTLA-4 antibody GIGA-564 induced less peripheral Treg proliferation, more efficient intratumoral Treg depletion, superior anti-tumor efficacy, and less toxicity in human *CTLA4* knock-in mouse models. This work suggests a translational path forward to improve outcomes for cancer patients.

## RESULTS

### *In vitro* characterization of a diverse panel of anti-CTLA-4 antibodies

We generated a library of natively paired single chain variable fragments (scFvs) specific to CTLA-4 by immunizing Trianni Mouse® animals, which produce antibodies with fully-human variable domains and mouse constant regions; we then used our proprietary microfluidics platform and yeast scFv display to select for CTLA-4 binders (39, 40, 41). Fourteen of the antibodies were cloned as full-length human IgG1 antibodies, produced in Chinese hamster ovary (CHO) cells, and were shown to bind to recombinant human CTLA-4 by surface plasmon resonance (SPR) with low nanomolar affinity (average equilibrium dissociation constant, KD: 8.7 nM), similar to that of ipilimumab (**Fig. S1A-B** and **Table S1**). These 14 mAbs bound CHO cells stably expressing human CTLA-4, with no off-target binding to CHO cells expressing the irrelevant target CD27 (**Fig. S1C** and **Table S1**). Next, the ability of these mAbs to block interaction of CTLA-4 with its endogenous ligands (the B7 proteins CD80 and CD86) was assessed by a cell-based assay. The mAbs exhibited a variety of half maximal effective concentration (EC50, µg/mL; average: 0.34; range: 0.1-2.17), and a broad range of maximum signals (average: 240; range: 13-548), where lower values indicated less blocking (**Fig. S1D** and **Table S1**). The correlations between affinity KD and blocking EC50 or between affinity KD and blocking maximum signal were not significant (linear regression; R^2^=0.06 and R^2^=0.16, respectively; p>0.05, **Fig. S1E**). For this set of 14 antibodies, we speculate that B7 ligand blocking is a function of anti-CTLA-4 binding epitope and not CTLA-4 binding affinity. Of interest, aCTLA-4.15, subsequently re-named GIGA-564, showed a combination of high affinity and low blocking maximum signal.

### FcR effector function is required for the anti-tumor efficacy of ipilimumab in a murine model

Fc receptor (FcR) binding and the corresponding depletion of intratumoral Tregs has been shown to be necessary for efficacy of anti-CTLA-4 mAbs in murine models (14, 15, 16, 17, 18, 19), however, the role of Treg depletion in the efficacy of CTLA-4 mAbs has remained controversial (42). Additionally, the role of the Fc effector function in the anti-tumor efficacy of ipilimumab has not been reported. Thus, we wished to investigate the anti-tumor effect of conventional anti-CTLA-4 mAbs, including ipilimumab, in the presence and absence of Fc effector functions.

First, we generated wild-type human IgG1 and human IgG1 N297Q mutant Fc variants of ipilimumab and another strong blocker, aCTLA-4.28 (**Fig. S1** and **Table S1**). The N297Q mutation eliminates Fc glycosylation and the ability to bind FcRs, thus abrogating Fc effector functions such as ADCC (43). The CTLA-4 mAbs with the N297Q mutation were still able to bind CTLA-4^+^ CHO cells (**Fig. S2**). As ipilimumab does not bind murine CTLA-4, a human *CTLA4* knock-in (hCTLA-4 KI) mouse model, where the human *CTLA4* coding region is knocked-in to the murine *Ctla4* locus, was used for *in vivo* efficacy studies. The majority (89.9% and 91.5%, respectively) of CD4 conventional T cells (CD4^+^FOXP3^-^) in the lymph node of hCTLA-4 KI and wildtype mice were naïve (CD44^Lo^CD62L^+^) (**Fig. 1A**), suggesting that CTLA-4 is functional in these hCTLA-4 KI mice. Both the ipilimumab analog and aCTLA-4.28 with a wildtype human IgG1 Fc led to strong regression of MC38 tumors (colon cancer model) in hCTLA-4 KI mice (**Fig. 1B**). However, in mice treated with ipilimumab_N297Q or aCTLA-4.28_N297 tumor growth was not significantly different from isotype control treated mice (**Fig. 1B**). These data indicate that anti-CTLA-4 mAbs, including ipilimumab, require FcR binding for anti-tumor activity and that blocking the binding of B7 ligands to CTLA-4 is not sufficient for the anti-tumor activity of anti-CTLA-4 mAbs.

**Fig. 1.**
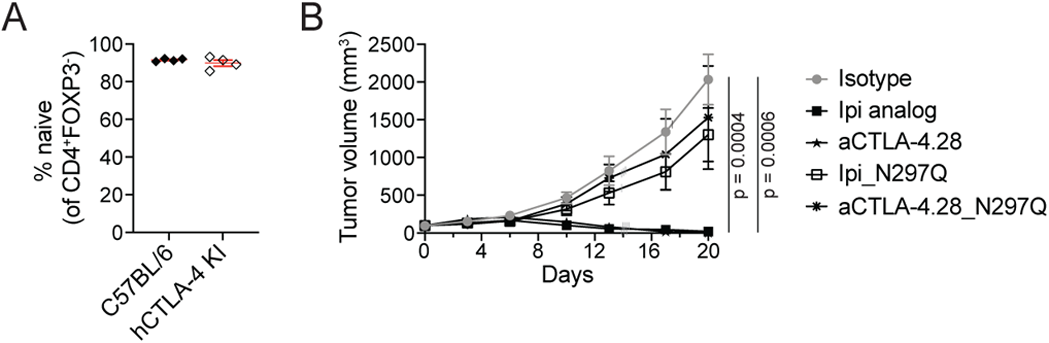
FcR effector function is required for the anti-tumor efficacy of ipilimumab in a murine model. **A**. The plot shows the frequency of CD4+FOXP3− T cells from the LN of individual control (C57BL/6) or hCTLA4-KI mice that are CD44LoCD62L+ as determined by flow cytometry. Each symbol represents data from individual animals (n=4 animals per a group) and red lines indicate means +/− SEM. **B.** hCTLA-4 KI mice bearing MC38 tumors were randomized when tumors were 50 – 151 mm3 (day 0) and treated bi-weekly with the indicated antibody at 5 mg/kg for 5 doses. The plot shows the tumor volume (mean ± SEM) of MC38 tumors over time that were treated with the indicated antibody. Tumor volume from mice euthanized due to tumor burden above 3000 mm3 was carried forward. Vertical lines indicate censored data: gray vertical lines indicate animals lost post dosing, likely due to anti-drug antibody (ADA)-induced hypersensitivity, black vertical lines refer to animals euthanized due to tumor burden and for which the data was carried forward. Ipi analog indicates an ipilimumab mAb produced and purified by us; N297Q indicates that the antibody contains an N297Q mutation in the Fc domain of the antibody to abrogate Fc effector function. n = 8 animals for isotype, aCTLA-4.28, and Ipi-N297Q; n=7 animals for ipi analog and aCTLA-4.28. p = 0.0004 when comparing Ipi analog to isotype and p = 0.0006 when comparing aCTLA-4.28 to isotype for change in tumor volume between groups (linear mixed effects model).

### GIGA-564 has weak ability to block CTLA-4 from binding to CD80/CD86

To confirm the weaker blocking ability of GIGA-564, we performed ELISAs to separately determine the ability of CD80 or CD86 to bind CTLA-4 in the presence of CTLA-4 mAbs (**Fig. 2A**). If binding of the mAb to CTLA-4 blocked CTLA-4 from binding to the ligand, the amount of ligand detected would decrease with increasing mAb concentration. GIGA-564 had much weaker blocking efficiency for both CD80 and CD86 than ipilimumab or aCTLA-4.28, as the curves were strongly shifted to the right (**Fig. 2B** and **Table S2**). Additionally, GIGA-564 was unable to fully block either ligand, even at the highest concentrations (**Fig. 2B**). This is consistent with its inability to induce full CD28 signaling in the cell-based signaling assay (**Fig. S1D**), which required the mAbs to strongly block binding of CTLA-4 to CD80 and CD86.

**Fig. 2.**
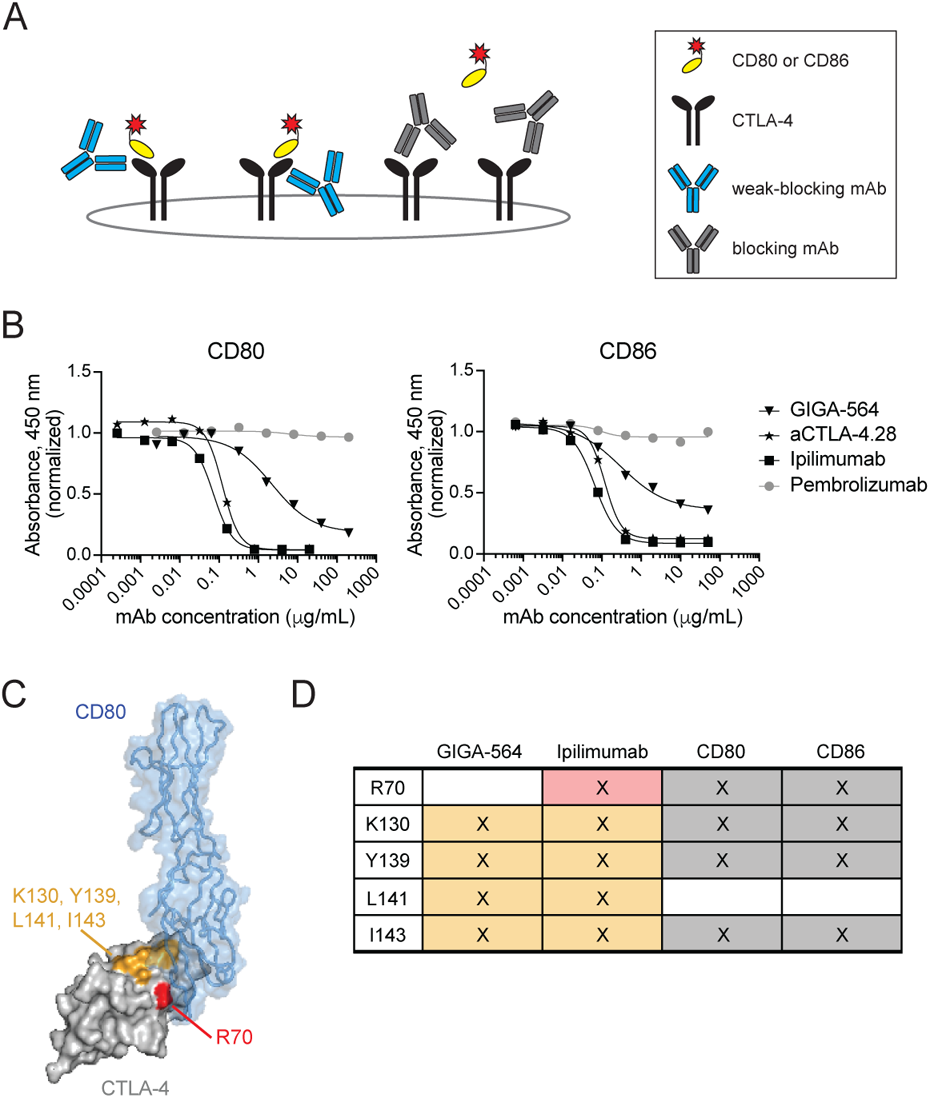
GIGA-564 has little ability to block the interaction of CTLA-4 and CD80/CD86 in vitro. **A.** The ability of CTLA-4 mAbs to block binding of CD80 or CD86 was measured using a plate-based ELISA method. CTLA-4 was used to coat the plate, then after the antibody samples were incubated, His-tagged CD80 or CD86 was added, and the amount of ligand able to bind to CTLA-4 was measured. Blocking mAbs that prevent CD80 or CD86 from binding to CTLA-4 reduce the absorbance signal due to lack of CD80 or CD86 binding to CTLA-4. Weak-blocking mAbs still allow CD80 or CD86 to bind, preventing the loss of all absorbance signal. **B.** Plots show the ability of GIGA-564 to block the interaction between CTLA-4 and the B7 ligands CD80 and CD86 compared to ipilimumab and CTLA-4.28, as assessed by ELISA as described in (A). Absorbance values were normalized to an anti-PD-1 control (pembrolizumab) and displayed as the average of two technical replicates. **C-D.** The key residues mediating CTLA-4 binding were identified for GIGA-564 and ipilimumab by shotgun mutagenesis of CTLA-4, followed by staining and flow cytometry assessment of binding. **C.** Shown is the crystal structure (Protein Database [PDB] 1I8L) of the complex between CD80 (blue) and CTLA-4 (gray) on which the CTLA-4 epitope residues shared between ipilimumab and GIGA-564 were colored orange and the key differentiating residue R70 was colored red (visualized with Pymol). Additionally, G142 was identified as a secondary residue for the epitope of ipilimumab but not GIGA-564. **D.** Table showing key amino acids on CTLA-4 of interest for these epitopes; those found by mutational analysis to be important for binding of CTLA-4 to CD80 or CD86 in a cell-based assay are marked in gray to indicate the epitope residues for those proteins (45).

The key amino acids in the CTLA-4 epitope for GIGA-564 and ipilimumab were determined using the shotgun mutagenesis epitope mapping method (44). Epitope mapping revealed that GIGA-564 and ipilimumab have overlapping but distinct epitopes. In particular K130, Y139, L141, and I143 are key amino acids that are part of the epitope for both GIGA-564 and ipilimumab (**Fig. 2C-D**). However, R70 is a key amino acid for the epitope of ipilimumab but not GIGA-564 (**Fig. 2C-D**). Importantly, R70 is also important for CD80 and CD86 binding to CTLA-4 (**Fig. 2D**) (45). We speculate that R70 on CTLA-4 may be more available to bind CD80 and CD86 when CTLA-4 is bound by GIGA-564 compared to ipilimumab, explaining why GIGA-564 only minimally blocks CD80/CD86 binding to CTLA-4.

### GIGA-564 treatment induces anti-tumor responses in murine models

We next tested the anti-tumor efficacy of GIGA-564. Both GIGA-564 and commercial ipilimumab led to nearly complete control of MC38 tumors in hCTLA-4 KI mice when they were dosed at 5 mg/kg twice a week (**Fig. 3A**). Similarly, in hCTLA-4 KI mice bearing RM-1 tumors (prostate cancer model), which are more resistant to anti-CTLA-4, GIGA-564 led to similar tumor growth inhibition to commercial ipilimumab when dosed at 5 mg/kg on days 0, 3, and 6 (**Fig. 3B**). Thus, we found that anti-CTLA-4 anti-tumor activity is independent of checkpoint inhibition via CTLA-4 blocking in multiple tumor models.

**Fig. 3.**
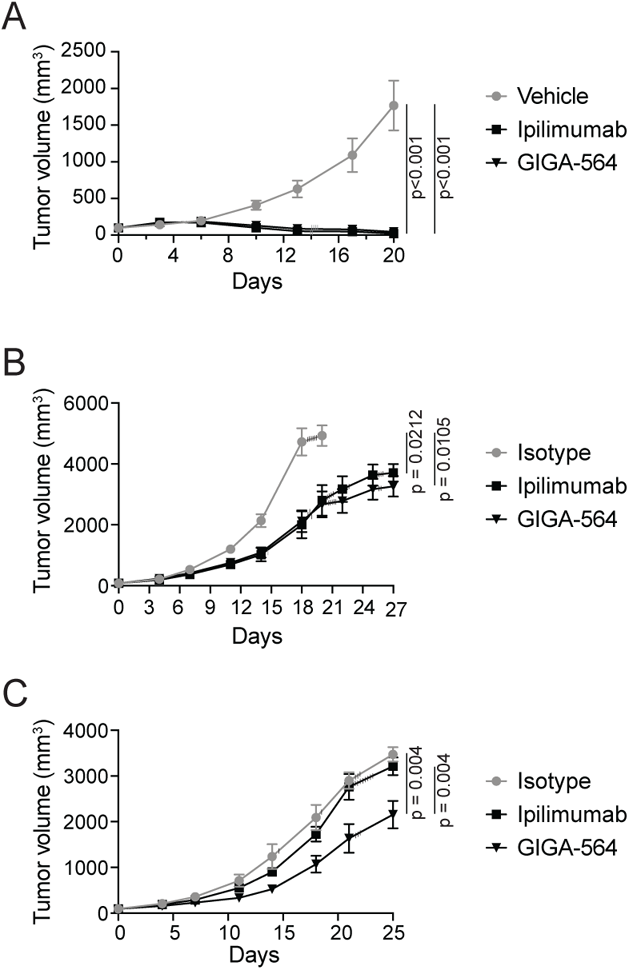
GIGA-564 inhibits tumor growth in murine models. **A.** hCTLA-4 KI mice bearing MC38 tumors were randomized when tumors were 50 – 150 mm3 (day 0) and treated bi-weekly with the indicated antibody at 5 mg/kg for 5 doses. Plots show the tumor volume (mean ± SEM) of MC38 tumors over time that were treated with the indicated antibody. Tumor volume from mice euthanized due to tumor burden above 3000 mm3 was carried forward. Gray vertical lines indicate censored data (animals lost post dosing likely due to ADA-induced hypersensitivity). n = 7 animals for GIGA-564 and n = 6 animals for vehicle and commercial ipilimumab. **B.** hCTLA-4 KI mice bearing RM-1 tumors were randomized when tumors were 40 – 125 mm3 (day 0) and treated on day 0, 3, and 6 with 5 mg/kg of the indicated antibody. Plots show the tumor volume (mean ± SEM) of RM-1 tumors over time that were treated with the indicated antibody. Tumor volume from mice euthanized due to tumor burden above 3000 mm3 was carried forward until no mice from that group were alive (black vertical lines). Vertical lines indicate censored data. The gray vertical line indicates one mouse from the ipilimumab treated group was euthanized due to tumor ulceration. n = 11 animals for ipilimumab and GIGA-564 treatment and n = 7 animals for isotype treatment. **C.** hCTLA-4 KI mice bearing MC38 tumors were randomized when tumors were 65 – 125 mm3 (day 0) and treated on days 0, 3, and 6 with 0.3 mg/kg of the indicated antibody. Plots show the tumor volume (mean ± SEM) of MC38 tumors over time that were treated with the indicated antibody. Tumor volume from mice euthanized due to tumor burden above 3000 mm3 was carried forward until no mice from that group remained alive (black vertical lines). n = 13 animals for ipilimumab and GIGA-564 and n = 8 animals for isotype treated groups. P-values are shown for statistically significant differences in changes in tumor volume measured longitudinally (linear mixed effects model).

Highly efficacious dosing of anti-CTLA-4 mAbs at 5 mg/kg made it difficult to identify any potential difference in the anti-tumor efficacy of ipilimumab and GIGA-564. Therefore, the ability of 0.3 mg/kg of ipilimumab or GIGA-564 to control progression of MC38 tumors in hCTLA-4 KI mice was assessed. GIGA-564 was more effective in limiting tumor progression than commercial ipilimumab at this low dose (**Fig. 3C**).

### GIGA-564 induces less peripheral Treg proliferation and efficiently depletes intratumoral Tregs

Co-stimulation is well known to be important for effector T cell proliferation, but it is also important for Treg proliferation (25, 26); for example, Tregs stimulated via CD3/CD80 proliferate more than Tregs stimulated via CD3 alone, and CTLA-4 blockade overcomes inhibition by exogenous CTLA-4-Fc (**Fig. S3)**. In agreement with this, blocking or acute loss of CTLA-4 enhances Treg proliferation *in vivo* (12, 16, 21, 22, 23). Thus, we hypothesized that GIGA-564 would induce less Treg proliferation than ipilimumab. To test this possibility, we treated hCTLA-4 KI mice bearing established MC38 tumors with mAbs at 5 mg/kg on days 0, 3, and 6 and analyzed the proliferation status of peripheral T cell subsets from a non-draining lymph node on day 7 (**Fig. 4A**). This analysis revealed that ipilimumab increased the proliferation of Tregs more than it increased the proliferation of conventional T cells (Tconv) or CD8 T cells (**Fig. 4A**). Furthermore, GIGA-564 induced significantly less peripheral Treg proliferation than ipilimumab (**Fig. 4A**; Wilcoxon rank sum test, p = <0.05). These data are consistent with the *in vitro* data that showed that GIGA-564 only weakly blocks the interaction of CTLA-4 with its B7 ligands (**Fig. 2B**, **Fig. S1D**, and **Table S1**).

**Fig. 4.**
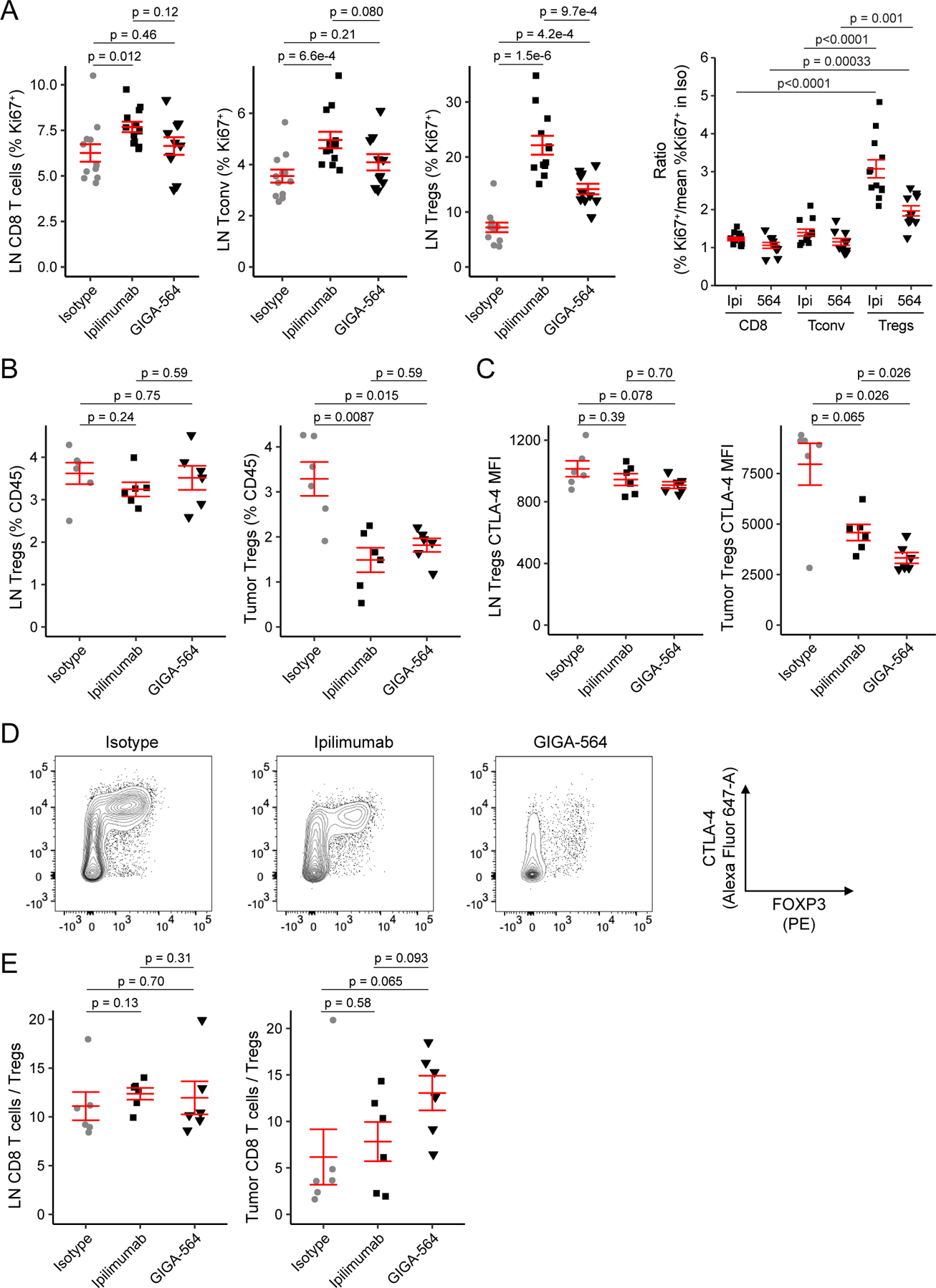
GIGA-564 induces less peripheral Treg proliferation but potently mediates depletion of intratumoral Tregs. **A.** hCTLA-4 KI mice (n = 12 animals) were treated with 5 mg/kg hIgG1 isotype control, ipilimumab, or GIGA-564 on days 0, 3, and 6, and euthanized for flow cytometry analysis on day 7. Two samples in the GIGA-564 group were excluded from analysis due to low cell count. The percent of CD8 T cells (Live, CD45+TCRβ+CD8+), CD4 Tconv (Live, CD45+TCRβ+CD4+FOXP3-), and Tregs (Live, CD45+TCRβ+CD4+FOXP3+) within the non-draining (left anterior axillary) lymph node (LN) expressing Ki67 was determined by flow cytometry (first 3 panels). The right panel shows the fold-change in the percent of cells of each subtype expressing Ki67 in ipilimumab or GIGA-564 treated mice relative to the mean frequency of that cell type expressing Ki67 in the isotype control treated group. **B-E.** hCTLA-4 KI mice bearing established MC38 tumors were randomized (n = 6 animals) and treated once with 5 mg/kg hIgG1 isotype control, ipilimumab, or GIGA-564 and cells from the non-draining LN and tumor were analyzed the following day by flow cytometry. **B.** Frequency of Tregs (as percentage of CD45+ cells) in LN (left) and tumor (right) in each treatment group. **C.** CTLA-4 geometric mean fluorescence intensity (MFI) in Tregs in LN (left) and tumor (right). **D.** Representative contour plots of CD4 T cells in each treatment group. X-axis and Y-axis correspond to FOXP3 and CTLA-4, respectively. **E.** Ratio of CD8 T cells relative to Tregs in LN (left) and tumor (right). P-values were calculated using a two-sided Wilcoxon rank sum test without adjustment for multiple comparisons.

Intratumoral Tregs express more surface CTLA-4 than effector T cells or even peripheral Tregs (14, 15, 16, 17). Thus, as FcR effector function activity of antibodies is dependent on the amount of antibody that binds to the cell surface (24), CTLA-4 mAbs selectively deplete intratumoral Tregs (15, 16, 17, 19, 21, 46). Furthermore, antibody-mediated depletion of CTLA-4^Hi^ Tregs in the tumor microenvironment (TME) has been shown to begin within the first 24 hours after antibody administration (16). Thus, to determine the ability of GIGA-564 to deplete intratumoral Tregs we treated hCTLA-4 KI mice bearing established MC38 tumors with 5 mg/kg of isotype, ipilimumab, or GIGA-564 and then analyzed Treg populations in the periphery and tumor by flow cytometry 1 day later. We found that both anti-CTLA-4 antibodies depleted intratumoral Tregs (Wilcoxon rank sum test, p<0.05) but not peripheral Tregs (**Fig. 4B**; Wilcoxon rank sum test, p>0.05). We also found that for animals administered GIGA-564, the population of remaining intratumoral Tregs had a lower CTLA-4 mean fluorescence intensity (MFI) than analogous populations from animals administered ipilimumab (**Fig. 4C-D**; Wilcoxon rank sum test, p<0.05). Additionally, GIGA-564 weakly enhanced the intratumoral CD8/Treg ratio compared to ipilimumab (**Fig. 4E**). These data suggest that GIGA-564 was more effective at depleting CTLA-4^Hi^ intratumoral Tregs, which in addition to the reduced Treg proliferation induced by GIGA-564, may explain why GIGA-564 is superior at controlling tumor growth at low doses than ipilimumab.

Additional flow analysis of the TME or peripheral lymph node of hCTLA-4 KI mice with established MC38 tumors treated with 2 or 3 doses of GIGA-564 or ipilimumab was performed (harvesting on day 4 or 7, respectively). Even with additional treatments, both GIGA-564 and ipilimumab were able to specifically deplete intratumoral Tregs but not peripheral Tregs (**Fig. S4**).

## GIGA-564 induces more FcR signaling than ipilimumab

The increased efficacy of low dose GIGA-564 compared to ipilimumab (**Fig. 3C**) may be in part due to the enhanced ability by which GIGA-564 depletes CTLA-4^Hi^ intratumoral Tregs (**Fig. 4C-E**). To investigate this, we compared the ability of GIGA-564 and ipilimumab to induce mouse FcR signal transduction. We found that GIGA-564 resulted in more FcR signaling than ipilimumab in Jurkat effector cells expressing murine FcγRIV, but not FcγRIII, when co-cultured with CHO cells stably expressing CTLA-4 (**Fig. 5A** and **Table S3**). This increased FcγRIV signaling induced by GIGA-564 may explain why GIGA-564 more efficiently depletes intratumoral Tregs than ipilimumab.

**Fig. 5.**
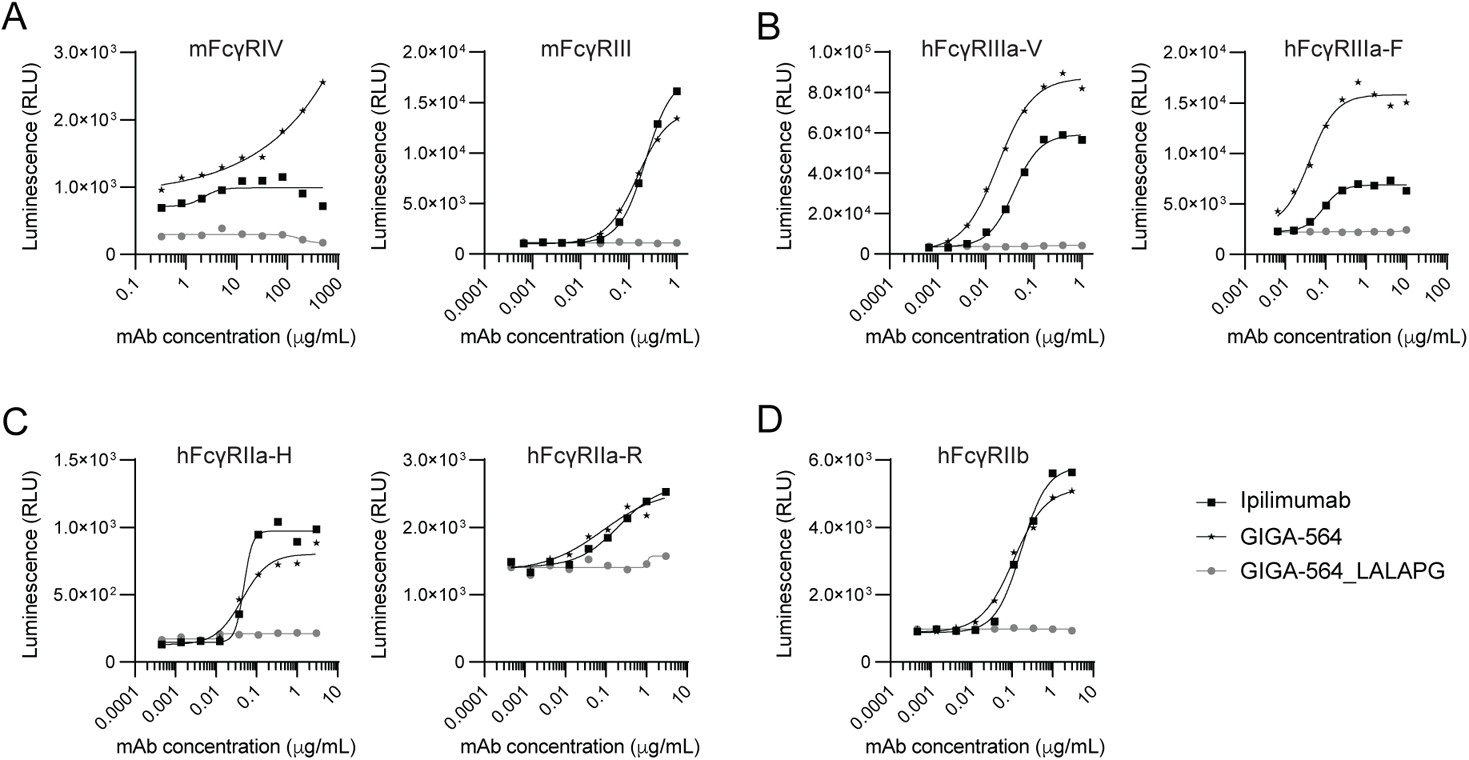
GIGA-564 induces more FcR signaling than ipilimumab. **A–D.** Target CHO cells expressing human CTLA-4 with the Y201G mutation to enhance surface expression were incubated with a titration series of ipilimumab (black squares), GIGA-564 (black stars), or a variant of GIGA-564 with LALA-PG mutations to disrupt FcγR binding (GIGA-564_LALA-PG, gray circles). Jurkat/NFAT-Luc effector cells with (A) mouse FcγRIV or FcγRIII, (B) human FcγRIIIa (high-affinity V158 or low-affinity F158 variant), (C) human FcγRIIa (high-affinity H131 or low-affinity R131 variant), or (D) human FcγRIIb were then added. Cells were incubated for 6 hours at 37°C then luciferase activity was measured. Data shown are relative luminescence units (RLU) emitting from the effectors cells and are plotted as the average of two technical replicates.

As the increased murine FcγRIV signaling induced by GIGA-564 may contribute to the enhanced depletion of intratumoral Tregs and efficacy induced by GIGA-564 in hCTLA-4 KI MC38 tumor-bearing mice, it was important to determine if GIGA-564 also induces more human FcR signal transduction. We found that GIGA-564 induced more human FcγRIIIa signaling than ipilimumab (**Fig. 5B** and **Table S3**). However, the increased FcR signaling induced by GIGA-564 was FcR-specific (**Fig. 5C-D** and **Table S3**). From a translational perspective, our data suggest that GIGA-564 may have increased ability to induce FcR signaling and thus FcR effector functions in patients, especially if expression of FcγRIIIa is used as a biomarker. The data further imply that even though some evidence suggests that ipilimumab has limited ability to deplete intratumoral Tregs in patients, GIGA-564 may act differently by inducing intratumoral Treg depletion and efficacy in patients.

It has previously been suggested that CTLA-4 mAbs with reduced binding at lower pH have enhanced ability to accumulate on the surface of Tregs because they dissociate from CTLA-4 in the early endosome. The hypothesis is that reduced binding at low pH allows for more CTLA-4 recycling, and thus more total CTLA-4 surface expression (47). However, we and others note that TMEs are often acidic and can have a pH below that of the early endosome (48). Thus, a CTLA-4 mAb with reduced binding at low pH would selectively bind CTLA-4 in the periphery over CTLA-4 in the TME. As our goal was to generate a CTLA-4 mAb that would target intratumoral Tregs, we determined the pH sensitivity of several CTLA-4 mAbs and found that GIGA-564 similarly bound to CTLA-4 at all tested pHs, including the at lower pHs often present in the TME (**Fig. S5A**).

FcR signaling can also be affected by several other factors including the amount of antibody that accumulates on the cell surface (24) and the affinity of the Fc portion of mAb to FcRs (43). We found no evidence that more GIGA-564 accumulates on the surface of CTLA-4^+^ cells than ipilimumab (**Fig. S5B**). The affinity of an Fc for FcγRIIIa is in part controlled by Fc glycosylation, specifically Fc fucosylation, because afucosylation enhances the affinity of the Fc for FcγRIIIa and thus ADCC (43). We found that GIGA-564 transiently expressed in ExpiCHO cells was less fucosylated than ipilimumab, while a CHO cell pool stably expressing GIGA-564 produced material that was 95% fucosylated (**Fig. S5C**). Human FcγRIIIa assays revealed that the amount of GIGA-564 afucosylation in each preparation correlated with FcγRIIIa signaling (**Fig. S5D**). However, further investigation revealed that GIGA-564 still demonstrates higher FcR activity than ipilimumab, even in the context of decreased afucosylation (**Fig. S5E**). The increased FcR activity of GIGA-564 likely contributes to its enhanced anti-tumor efficacy, and this may translate well to patients as GIGA-564 also induces enhanced human FcγRIIIa signaling.

### GIGA-564 induces durable anti-tumor immune responses

While first generation anti-CTLA-4s led to durable anti-tumor immunity (1, 2), little is known about the ability of an anti-CTLA-4 with minimal blocking activity to induce a durable anti-tumor immune response. To investigate this translationally important question, hCTLA-4 KI mice bearing established MC38 tumors were treated on days 0, 3, and 6 with control (isotype), ipilimumab, or GIGA-564 at 1 mg/kg, an FDA-approved dose of ipilimumab that induced a durable complete response in approximately 50% of animals (5/8 ipilimumab treated animals and 5/9 GIGA-564 treated animals) (**Fig. 6A**). Animals with a complete response on day 43 were then re-challenged with MC38 tumor cells on the opposite flank from the original tumor site. All durable complete responders in the group previously treated with commercial ipilimumab and 4/5 of the durable complete responders in the group previously treated with GIGA-564 did not develop tumors at the re-challenge site (**Fig. 6B**). Thus, GIGA-564, an anti-CTLA-4 with minimal blocking activity, leads to durable anti-tumor immunity by preventing tumor development at a distant site.

**Fig. 6.**
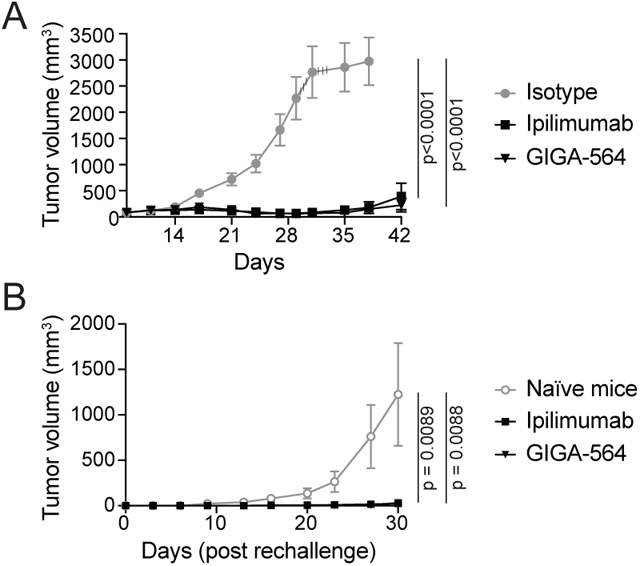
GIGA-564 provides protection in a murine tumor re-challenge model. **A.** hCTLA-4 KI mice bearing MC38 tumors were randomized when tumors were 60 – 120 mm3 (day 8) and then treated every 3 days for 3 doses with the indicated antibody at 1 mg/kg. Plots show the tumor volume (mean ± SEM) of MC38 tumors over time that were treated with the indicated antibody. Tumor volume from mice euthanized due to tumor burden above 3000 mm3 was carried forward (vertical black lines). n = 8 animals for isotype and commercial ipilimumab, n = 9 animals for GIGA-564. B. MC38 cells were implanted on the opposite flank of naïve C57BL/6 mice and mice from (A) that were previously treated with ipilimumab or GIGA-564 and had a durable complete response on day 43 (0 mm3 tumor volume). Plots show the tumor volume (mean ± SEM) of MC38 tumors over time that were previously treated with the indicated antibody (n = 5 animals previously treated with ipilimumab in [A] and 5 animals previously treated with GIGA-564 in [A]) or naïve mice (n = 5 animals). Tumor volume (in mm3) was analyzed using a linear mixed effects model including treatment group and day as fixed effects and animal identifier (ID) as a random effect to account for repeated measures. Statistical comparisons were made using the two-sided Wald test against the isotype control group in the initial challenge model and against the naïve control group in the re-challenge model. Compared to the naïve group, previous treatment with ipilimumab (p = 0.0089) or GIGA-564 (p = 0.0088) limited tumor growth.

### GIGA-564 induces less toxicity than ipilimumab in murine models

In the clinic, CTLA-4 antibodies are most frequently used in combination with PD-1 antibodies, but this combination can induce relatively high frequency of severe adverse events (49). Thus, a CTLA-4 mAb with enhanced efficacy and reduced toxicity would be beneficial to patients. Several lines of evidence reveal that a significant portion of the toxicities induced by conventional CTLA-4 mAbs likely arises through blocking CTLA-4 from binding CD80 and CD86 (30, 31, 32, 33, 34, 35, 36, 37, 38). This suggests that a CTLA-4 mAb with minimal blocking activity would not only have enhanced anti-tumor efficacy but may also result in less toxicity. Accordingly, we wanted to compare the toxicity induced by ipilimumab and GIGA-564 in the presence of anti-PD-1, as anti-CTLA-4 antibodies are most commonly used in combination with anti-PD-1 therapy.

hCTLA-4 KI mice treated with a CTLA-4 mAb have been shown to recapitulate some of the toxicity induced by CTLA-4 mAbs in patients (50), and repeated treatment of BALB/c mice with anti-mouse CTLA-4 antibodies was shown to induce intestinal inflammation (51). Thus, we treated hCTLA-4/hPD-1 double knock-in mice on the BALB/c background with vehicle, pembrolizumab, pembrolizumab plus ipilimumab, or pembrolizumab plus GIGA-564 every 3 days for 9 doses (**Fig. 7A**). One week after the last dose, mice were euthanized and tissues were collected for pathology analysis (**Fig. 7B-C** and **Fig. S6A-D**). Histopathological analysis revealed that pembrolizumab plus ipilimumab, but not pembrolizumab plus GIGA-564, induced more colonic epithelial damage and skin inflammation than vehicle treated mice or mice treated with pembrolizumab alone (**Fig. 7B-C**), while both combination treatments led to a minor increase in immune cell infiltration to the heart (**Fig. S6C-D**). As CTLA-4 mAbs have been shown to induce kidney deposition in hCTLA-4 KI mice on the C57BL/6 background, kidneys were collected on day 20 post initiation of treatment from hCTLA-4 KI mice bearing MC38 tumors treated with vehicle, 5 mg/kg ipilimumab, or 5 mg/kg GIGA-564 twice a week for 5 doses (**Fig. 3A**). Kidneys were processed into formalin fixed paraffin embedded (FFPE) blocks. FFPE sections were stained for anti-mouse immunoglobin and scored by a board-certified veterinary pathologist blinded from the study. Ipilimumab, but not GIGA-564, significantly increased the percent of glomeruli with murine Ig deposition (**Fig. S6E**), suggesting that ipilimumab induces more kidney damage than GIGA-564 in hCTLA-4 KI mice. Thus, GIGA-564 induces less toxicity than ipilimumab in these murine models.

**Fig. 7.**
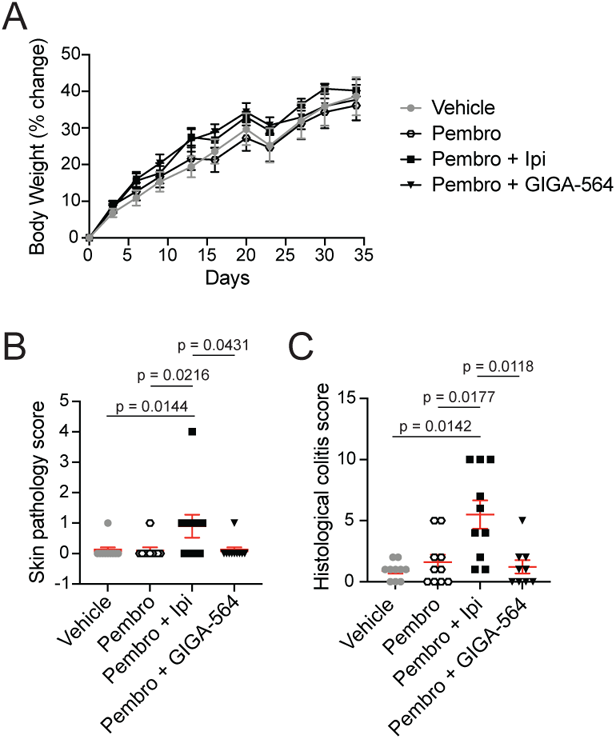
GIGA-564 plus pembrolizumab induces less toxicity than ipilimumab plus pembrolizumab in a murine model. hCTLA-4/hPD-1 double KI mice on the BALB/c background between 4-5 weeks of age were treated with vehicle, pembrolizumab, pembrolizumab plus ipilimumab (Ipi), or pembrolizumab plus GIGA-564 every 3 days for 9 doses. One week after the last dose mice were euthanized and tissues collected for pathology analysis. A. Plot shows percent change in body weight over time in mice treated with the indicated therapy (mean ± SEM, n = 10 animals). Four mice from the pembrolizumab plus ipilimumab group died on day 12, and from the pembrolizumab plus GIGA-564 treated group three mice died on day 12 and one mouse died on day 15, all likely due to post dosing ADA-induced hypersensitivity. A mixed effect model found no statistical differences in the percent body weight change between groups. B-C. Plots show skin inflammation (B) or colonic epithelial damage (colitis; C) scores induced by each treatment regimen. Adjusted p-values were calculated using the two-sided Dunn’s test applying the Benjamini-Hochberg step-down procedure to control for multiple comparisons.

In conclusion, GIGA-564 is a novel CTLA-4 mAb with minimal B7 ligand blocking activity resulting in enhanced efficacy but reduced toxicity in multiple murine models (**Fig. 8**), suggesting that reducing CTLA-4 checkpoint inhibition yields translationally relevant benefits for both efficacy and safety.

**Fig. 8.**
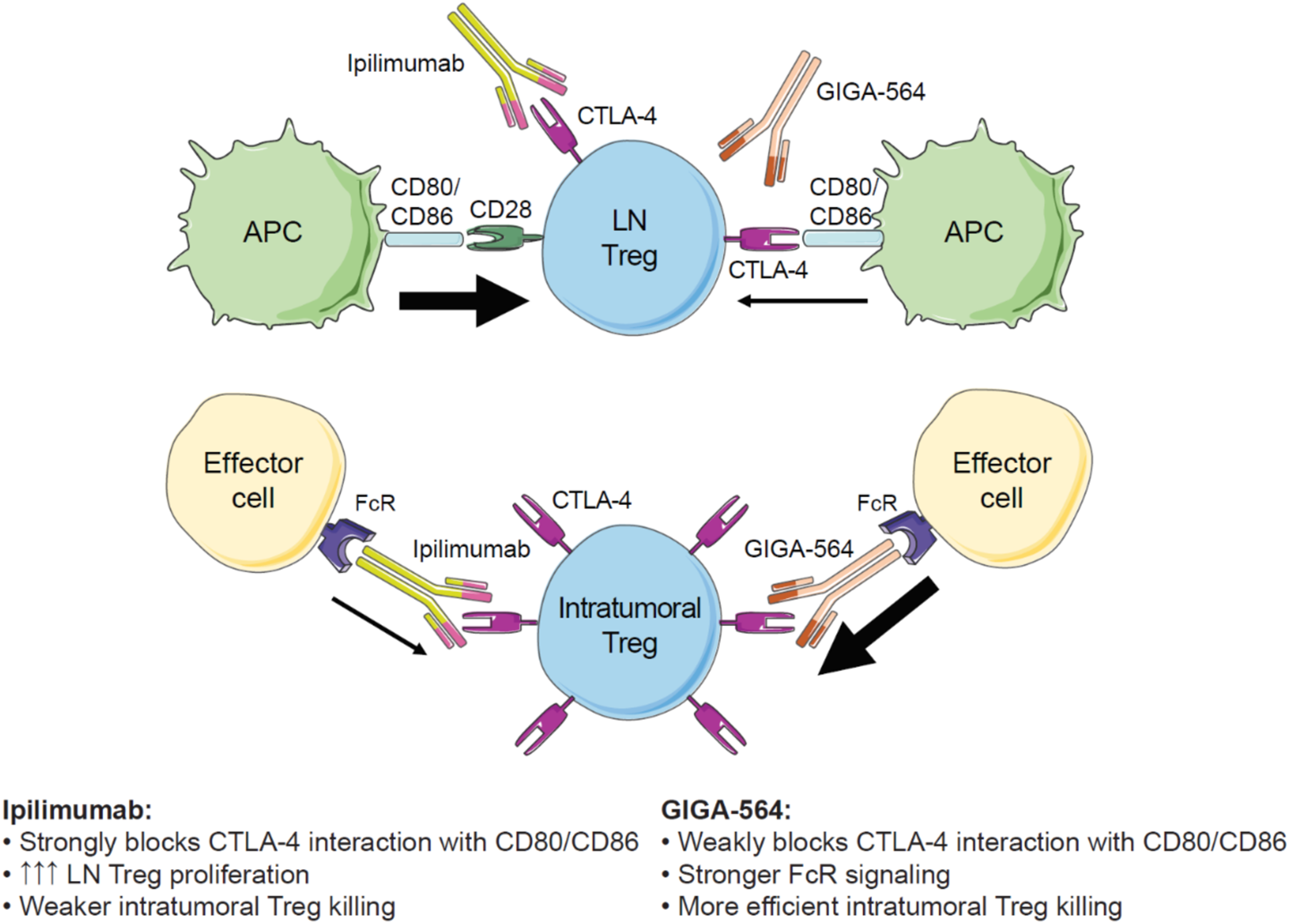
Model depicting ipilimumab and GIGA-564 mechanisms of action. Top panel: Ipilimumab blocks CTLA-4 interaction with CD80/CD86, which allows antigen presenting cells (APCs) to co-stimulate peripheral Tregs enhancing their proliferation. GIGA-564 weakly blocks CTLA-4 interaction with CD80/CD86 and thus induces less Treg proliferation. Bottom panel: Ipilimumab and GIGA-564 bind CTLA-4 on intratumoral Tregs to induce Treg killing via interactions with Fc receptor (FcR) on effector cells. GIGA-564 induces stronger FcR signaling and thus more efficiently depletes intratumoral Tregs than ipilimumab.

## DISCUSSION

Here we show that GIGA-564, a third generation anti-CTLA-4 with minimal ability to block CTLA-4 binding to its CD80/CD86 ligands and enhanced FcR activity, has superior anti-tumor activity and reduced toxicity compared to ipilimumab in murine models expressing human CTLA-4. A rationale for testing a CTLA-4 mAb with little ability to block CTLA-4 from binding CD80/CD86 was that CTLA-4 is more highly expressed by Tregs than effector cells (15) and Treg proliferation is also enhanced by co-stimulation (25, 26), suggesting that blocking CTLA-4 binding to CD80/CD86 may preferentially enhance proliferation of Tregs. Indeed, we found that in hCTLA-4 KI mice, ipilimumab enhanced Treg proliferation more than proliferation of CD4^+^FOXP3^-^ or CD8 T cells. This result is analogous to clinical and pre-clinical data for first-generation IL-2 therapy, which also preferentially enhanced Tregs over effector T cells (27, 28, 29). Second generation IL-2 therapies that do not bind CD25 on Tregs resulted in less Treg proliferation and have had promising results in early clinical trials (29, 52). Similarly, GIGA-564 leads to less Treg proliferation and increased anti-tumor activity in a murine model when compared to the first generation anti-CTLA-4 mAb ipilimumab.

It has recently been suggested that anti-CTLA-4-mediated depletion of intratumoral Tregs may be transient (42), explaining why intratumoral Treg depletion is not obvious at all timepoints following anti-CTLA-4 therapy (53). Thus, one possibility is that anti-CTLA-4 mediated expansion of peripheral Tregs provides a reservoir that reseeds the intratumoral Treg niche. Additionally, while anti-CTLA-4 mAbs efficiently induce depletion of intratumoral Tregs in the MC38 model primarily used here, the ability of anti-CTLA-4s to deplete intratumoral Tregs in patients is expected to be variable. This is because the expression of CTLA-4 by intratumoral Tregs varies among and within patients (15) as can the affinity of FcRs (17, 54), which are able to mediate antibody-dependent killing of CTLA-4^Hi^ Tregs (15, 55). Thus, in some patients, CTLA-4 mAbs that block CD80/CD86 binding to CTLA-4 may actually enhance intratumoral Treg levels. It has recently been shown that anti-PD-1 blockade can also enhance proliferation of Tregs in patients with PD-1^Hi^ intratumoral Tregs, leading to reduced efficacy and even hyperprogression (56, 57). Taken together, these data suggest that patients may benefit from clinical translation of an anti-CTLA-4 with reduced ability to enhance Treg proliferation. Furthermore, GIGA-564 may synergize well with anti-PD-1s, because it would be expected to deplete the intratumoral Tregs an anti-PD-1 could induce.

Another rationale for identifying and testing a CTLA-4 mAb with little ability to block CTLA-4 binding to its CD80/CD86 ligands is that several lines of evidence suggest that blocking the ability of CTLA-4 to bind CD80/CD86 may strongly contribute to the severe adverse events induced by first generation CTLA-4 mAbs. First, *CTLA4* mutations in humans that decrease binding of CTLA-4 to its B7 ligands are associated with immune infiltration into the brain, gastrointestinal tract, and lung, and can result in diarrhea, colitis, lymphocytic interstitial lung disease, and occasionally autoimmune thyroiditis (30, 31, 32, 33, 34, 35, 36). Clinically it has also been reported that the gastrointestinal biopsies of CHAI patients (<u>C</u>TLA-4 <u>h</u>aploinsufficiency with <u>a</u>utoimmune infiltration) are reminiscent of patients treated with anti-CTLA-4 blocking antibodies (30, 32). Other relevant clinical data has been reported after treatment with abatacept (CTLA-4-Ig), which strongly binds the B7 proteins CD80 and CD86 (31, 33). At least one patient treated with abatacept had greatly reduced gastrointestinal symptoms, including diarrhea (31). Additionally, abatacept “has achieved substantial and lasting improvement of interstitial lung disease in LATAIE patients” (32). Taken together, these data suggest that in humans, the loss of CTLA-4 binding to its B7 ligands induces immune infiltration into non-immune organs and pathology similar to that induced by blocking CTLA-4 mAbs. It has also recently been shown that checkpoint inhibitor-induced colitis in patients is not associated with a loss of Tregs and thus may be caused by CTLA-4 blockade rather than depletion of CTLA-4^Hi^ Tregs (38). In agreement with this, our current study found that pembrolizumab plus GIGA-564 induced less colonic epithelial damage and skin pathology than pembrolizumab plus ipilimumab in human CTLA-4/PD-1 double knock-in mice on the BALB/c background. Gastrointestinal and dermatological adverse events in patients are among the most common severe adverse events associated with ipilimumab or ipilimumab plus anti-PD-1 therapy (58, 49). Thus, our data suggest that clinical translation of GIGA-564 may result in reduced rates of these adverse events commonly associated with conventional CTLA-4 mAbs and thus benefit patients.

Surprisingly, we found that GIGA-564 has enhanced FcR activity *in vitro* and *in vivo*. As next-generation anti-CTLA-4s with enhanced FcR activity have improved anti-tumor efficacy (19), it is likely that the enhanced FcR activity of GIGA-564 contributes to the improved anti-tumor efficacy of this mAb. Our data indicate that the enhanced FcR function of transiently produced GIGA-564 may be due to the partially afucosylated status of this material; however, further investigation revealed that GIGA-564 still demonstrated higher human FcγRIIIa activity even in the context of decreased afucosylation, suggesting that our results will translate to patients. Of note, the GIGA-564 material used in the murine toxicity study had relatively high levels of afucosylation (80.9%) and Fc effector function, further emphasizing the role of checkpoint inhibition in the induction of adverse events induced by conventional anti-CTLA-4 mAbs.

Another strategy that has been reported to enhance FcR function of CTLA-4 mAbs is selection of mAbs that dissociate from CTLA-4 at acidic pHs, leading to enhanced recycling of CTLA-4 and thus accumulation of the anti-CTLA-4 mAb on the cell surface (47). However, the tumor microenvironment is often acidic (48), and thus anti-CTLA-4s that preferentially bind at neutral pH will be less able to bind CTLA-4 in the tumor microenvironment, precisely where we aim to selectively deplete Tregs. Importantly, GIGA-564 was able to bind to CTLA-4 efficiently at low pH and can thus target intratumoral Tregs.

It has been argued that the FcR-dependent activity of anti-CTLA-4s may result from FcRs holding CTLA-4 away from the immune synapse and thus serves to further functionally inhibit the CTLA-4/B7 immune checkpoint (59). As CTLA-4 is most highly expressed by Tregs and co-stimulation is also important for Treg proliferation (15, 25, 26), anti-CTLA-4s that block the ability of CTLA-4 to bind its B7 ligands induce significant Treg proliferation. Thus, the fact that GIGA-564 only weakly blocks the ability of CTLA-4 to bind its B7 ligands and has enhanced FcR activity but induces less Treg proliferation than ipilimumab, suggests that the Fc activity of anti-CTLA-4s play little role in the inhibiting the ability of anti-CTLA-4s to bind its B7 ligands.

In conclusion, we showed that the weak blocking CTLA-4 mAb GIGA-564 induced less proliferation of peripheral Tregs, more efficiently depleted intratumoral CTLA-4^Hi^, and had superior anti-tumor activity while inducing less toxicity than ipilimumab in murine models expressing human CTLA-4. In the near future we expect to move GIGA-564 toward large-scale GMP manufacturing and regulatory approval for first-in-human studies in patients with life-threatening cancers.

## MATERIALS AND METHODS

### Antibody sequences

Amino acid sequences for the IgK and IgG variable regions for the 14 antibodies described here that were expressed as full-length antibodies and tested for *in vitro* blocking activity are listed in **Table S4**.

### Generation of full-length antibodies and expression in CHO cells

Five Trianni Mouse® mice expressing antibodies with fully-human variable regions were immunized at Antibody Solutions (Sunnyvale, CA, USA), as described elsewhere (41). Briefly, mice were immunized with soluble His-tagged CTLA-4 (CT4-H5229; Acro Biosystems, Newark, DE, USA) and alendronate (ALD) / muramyl dipeptide (MDP) adjuvant twice per week for four weeks. Mice were euthanized and inguinal and popliteal lymph nodes and spleen were harvested and processed into a single cell suspension. Cells from all mice were pooled by tissue type and B cells were selected from lymph node and spleen using a mouse Pan-B negative selection kit (Stemcell Technologies, Vancouver, BC, Canada). Generation of natively paired heavy and light chain libraries, yeast surface display of scFvs and FACS sorting, and antibody repertoire analysis was described elsewhere (41).

From this analysis, scFv sequences were selected for full-length antibody expression based on enrichment after sorting. Expression constructs were Gibson assembled using GeneBlocks (Integrated DNA Technologies, Coralvile, IA, USA) and NEBuilder HiFi DNA Assembly Master Mix (NEB, Ipswich, MA, USA) to integrate sequences into a vector appropriate for transient expression in CHO cells. The vector used was a variant of the pCDNA5/FRT mammalian expression vector (Thermo Fisher Scientific, Waltham, MA, USA). The vector has an elongation factor 1 alpha (EF1α) promoter to express light chain followed by a bovine growth hormone (BGH) polyA sequence and a cytomegalovirus (CMV) promoter to express heavy chain followed by a second BGH polyA sequence. All constructs were synthesized as human IgG1 isotype, regardless of a given antibody’s IgG isotype in the original repertoire. Constructs were transformed into NEB 10-beta *E. coli* for amplification and purified with the ZymoPURE Plasmid Maxiprep Kit (Zymo Research, Irvine, CA, USA). The purified plasmid was then used for transient transfection in the ExpiCHO system (Thermo Fisher Scientific, Waltham, MA, USA). Transfected cells were cultured for 7–9 days in ExpiCHO medium, and then antibodies were purified from filtered supernatant using Protein A columns (MilliporeSigma, St. Louis, MO, USA). Antibody purity and proper size was verified by Coomassie stained sodium dodecyl sulphate-polyacrylamide gel electrophoresis (SDS-PAGE) (Thermo Fisher Scientific, Waltham, MA, USA).

### Transient transfection and mAb protein production

For larger-scale expression, the full-length kappa chain of GIGA-564 was cloned into a separate pSF expression vector (MilliporeSigma, St. Louis, MO, USA) with a CMV promoter. The immunoglobulin kappa-only and dual-gene (immunoglobulin kappa and immunoglobulin gamma) plasmids were transfected at a 2:1 molar ratio into ExpiCHO cells using ExpiFectamine (Thermo Fisher Scientific, Waltham, MA, USA). Briefly, for every 100 mL of culture 100 μg of total plasmid was mixed with 320 μl of ExpiFectamine in 4 mL of OptiPro serum free media (Thermo Fisher Scientific, Waltham, MA, USA), incubated at room temperature for 5 minutes, then added to cells freshly passaged to 6 × 10^6^ cells/mL. Cells were fed on day 1 and day 5 after transfection with 16 mL of ExpiCHO feed, then harvested on Day 8 or when the viability dropped below 75%.

Harvested cell-culture fluid (HCCF) was loaded onto a HiTrap MabSelect Prism A column (Cytiva, Marlborough, MA, USA), equilibrated with PBS pH 7.0-7.4 using a fast protein liquid chromatography (FPLC) instrument (AKTA pure 25, Cytiva, Marlborough, MA, USA), and eluted with 100 mM citrate pH 3. Fractions were pooled to collect >90% of eluted material and neutralized to pH 6.2 using 1 M tris pH 9. Neutralized eluate was dialyzed into 40 mM histidine + 240 mM sucrose pH 6.2 and formulated with 0.2% tween-20. Concentration was determined by absorbance (NanoDrop 8000, Thermo Fisher Scientific, Waltham, MA, USA) and endotoxin quantified by limulus amebocyte lysate assay (NexGen PTS, Charles River, Wilmington, MA, USA). Routine biophysical characterization included size exclusion chromatography high performance liquid chromatography (SEC-HPLC; 7.8 × 300 mm, 2.7 μM, 300A column, Agilent, Santa Clara, CA, USA), SDS-PAGE (12% tris-glycine, Thermo Fisher Scientific, Waltham, MA, USA), and capillary electrophoresis sodium dodecyl sulphate (CE-SDS; protein 230, BioAnalyzer 2100, Agilent, Santa Clara, CA, USA).

### Clonal cluster analysis and visualization

We used USEARCH (60) to compute the total amino acid differences between each pairwise alignment of FACS-sorted scFv sequences. We then used the R package igraph (version 1.2.6) (61) to generate a clustering plot for the pairwise alignments. The sequences were represented as “nodes”, while “edges” were the links between nodes. Edges indicate pairwise alignments with ≤9 amino acid differences. The layout_with_graphopt (charge = 0.03, niter = 1000) option was used to format the output.

### Affinity measurements

Kinetic analysis of CTLA-4 mAb HCCF from CHO expression was performed on an MX-96 instrument (IBIS Technologies, Enschede, Netherlands) by Carterra (Dublin, CA, USA). A medium density capture chip was created with anti-Human IgG Fc (SouthernBiotech, Birmingham, AL, USA) with a surface density of 925-1200 RU. A CFM printer (IBIS Technologies, Enschede, Netherlands) printed one 10 minute print of mAb HCCF for each sample. His-tagged human CTLA-4 antigen (CT4-H5229; Acro Biosystems, Newark, DE, USA) injections for kinetic analysis were 5 minutes and dissociations were 10 minutes. Kinetic analysis was performed with five 5-fold serial titrations starting at 500 nM antigen and fitted to a 1:1 monovalent model.

### Flow cytometry for in vitro characterization of mAbs

To determine if the CTLA-4 mAbs bound the appropriate antigen expressed on the surface of mammalian cells, CHO cell lines stably expressing human CTLA-4 or an irrelevant antigen (CD27) were generated. Cells (0.5 × 10^6^ each cell line) were combined and washed with MACS buffer (Dulbecco’s Phosphate Buffered Saline [DPBS], with 0.5% bovine serum albumin [BSA], and 2mM ethylenediaminetetraacetic acid [EDTA]). The cells were incubated with 10 μg/mL anti-CTLA-4 for 30 min at 4°C, then excess, unbound mAbs were removed by washing twice with MACS buffer. The cells were then stained with PE-conjugated anti-Human IgG Fc antibody (clone M1310G05, BioLegend 410720, San Diego, CA, USA) to detect bound anti-CTLA-4 and with fluorescein isothiocyanate (FITC)-conjugated anti-CD27 (clone O323,

BioLegend 302806, San Diego, CA, USA) to distinguish the cells expressing the irrelevant antigen. Cells were washed twice with MACS buffer, fixed with 4% paraformaldehyde fixation buffer (BioLegend 420801, San Diego, CA, USA) for 20 minutes at room temperature and washed twice more with MACS buffer. For validation of the N297Q variants of the ipilimumab analog and aCTLA-4.28, a similar procedure was followed except CHO cells lacking expression of human CTLA-4 were used in place of the CD27-expressing cells, the two cell lines were stained separately, FITC-conjugated anti-Human IgG Fc antibody (clone M1310G05, BioLegend 410719, San Diego, CA, USA) was used to detect bound anti-CTLA-4, cells were not treated with fixation buffer, and cells were instead stained with 4′,6-diamidino-2-phenylindole (DAPI; BioLegend, San Diego, CA, USA) and gated for live (DAPI^-^) cells on the cytometer. Binding of mAbs to CTLA-4 on the cell surface was determined using LSR II (BD Biosciences, San Jose, CA, USA) or CytoFLEX LX (Beckman Coulter, Brea, CA, USA) flow cytometers and analyzed using FlowJo (v10.6.1, BD Biosciences, San Jose, CA, USA).

To determine accumulation of anti-CTLA-4s on the cell surface, suspension CHO cells expressing wildtype human CTLA-4 were plated at 1 × 10^6^ cells/well in DPBS (Lonza, Basel, Switzerland) containing 0.5% BSA (MilliporeSigma, St. Louis, MO, USA) and 0.5 M EDTA (MilliporeSigma, St. Louis, MO, USA). Titrations of ipilimumab or GIGA-564 from 0-50 μg/mL were added to the cells and the plate was incubated at 37°C for 30 minutes to allow for internalization to occur. Anti-human IgG Fc allophycocyanin (APC) (BioLegend, San Diego, CA, USA) was added as a secondary antibody at 10 μg/mL and the plate was incubated at 4°C for 30 minutes. Following viability staining with DAPI (BioLegend, San Diego, CA, USA), data was acquired on the CytoFLEX LX (Beckman Coulter, Brea, CA, USA) and analyzed using FlowJo (v10.6.1, BD Biosciences, San Jose, CA, USA).

### Cell-based CTLA-4 blocking assay

CTLA-4 Blockade Bioassay was purchased from Promega (JA3001; Madison, WI, USA) and performed according to the manufacturer’s recommendations. Briefly, a serial dilution of each antibody was generated in assay buffer (90% RPMI 1640/10% FBS, supplied in the kit) in a sterile 96-well plate. CTLA-4 effector cells (provided with kit) were thawed and diluted into 3.2 mL of assay buffer and 25 μL of the cell suspension was added to each of the inner 60 wells of a 96-well, white flat-bottom plates. 25 μL of the appropriate antibody dilutions were added to the wells containing CTLA-4 effector cells. Artificial antigen presenting (aAPC)/Raji cells (provided with kit) were thawed and diluted into 7.2 mL of assay buffer, and 25 μL of the cell suspension was added to wells containing the diluted antibodies and CTLA-4 effector cells. Plates were incubated for 6 hours at 37°C in a tissue culture incubator with 5% CO_2_ before 75 μL of Bio-Glo Reagent was added to wells containing cell and antibody mixtures. Plates were incubated for 5-15 minutes at room temperature and luminescence was measured on a Spectramax i3x plate reader (Molecular Devices, San Jose, CA, USA). Data analysis was performed using the Softmax Pro (Molecular Devices, San Jose, CA, USA) or Prism (GraphPad, San Diego, CA, USA) software packages.

### CD80/CD86 blocking ELISAs

ELISA plates (Nunc, MaxiSorp ELISA plates, flat bottom, uncoated, BioLegend, San Diego, CA, USA) were coated with recombinant human CTLA-4-Fc (7268-CT, R&D Systems, Minneapolis, MN, USA) at 1 μg/mL overnight at 4°C. Plates were then blocked with 5% milk in phosphate buffered saline with tween 20 (PBST) for 1 hour at room temperature. A titration series of the indicated mAbs was then added to the plates and then the plates were incubated for 1 hour at room temperature to allow mAb binding. Excess, unbound mAb was removed by washing with PBST. Recombinant human His-tagged CD80 (R&D Systems 9050-B1, Minneapolis, MN, USA) or CD86 (R&D Systems 9090-B2, Minneapolis, MN, USA) was then added to the plates at 1 μg/mL. After incubation for 1 hour at room temperature, the plates were washed to remove unbound ligand. Bound ligand was detected with a horseradish peroxidase (HRP)-conjugated anti-His antibody (652504, BioLegend, San Diego, CA, USA). After incubation for 1 hour at room temperature and washing, the plates were developed with 3,3′,5,5′-Tetramethylbenzidine (TMB) substrate (34028, Pierce, Waltham, MA, USA). After sufficient signal was achieved, development was stopped by the addition of 1 N hydrochloric acid. Absorbance at 450 nm was read using a Spectramax i3x plate reader (Molecular Devices, San Jose, CA, USA). Half maximal inhibitory concentration (IC50) values were calculated by plotting absorbance versus the log of concentration using Prism (GraphPad, San Diego, CA, USA).

### Murine models

Murine experiments were done in compliance with all relevant ethical regulations and approved by the Institutional Animal Care and Use Committee of Crown Bioscience. Experiments using full-length hCTLA-4 knock-in HuGEMM^TM^ mice (Shanghai Model Organisms Center, Inc.) were performed at Crown Bioscience (Taicang Jiangsu Province, China). Full-length hCTLA-4 knock-in HuGEMM^TM^ mice were generated by knocking in human *CTLA4* cDNA with a polyA sequence at exon 1 of the murine *Ctla4* locus, replacing murine *Ctla4* expression with human *CTLA4* expression. Crown Bioscience acquired MC38 cells from FDCC (The Institutes of Biomedical Sciences (IBS), Fudan University, China) and RM-1 cells from SIBS (Shanghai Institutes for Biological Sciences, China) and authenticated cell line identity of research cell banks by single nucleotide polymorphism (SNP) analysis. Cell lines were mycoplasma negative. 8-12-week-old, female, full-length hCTLA-4 knock-in HuGEMM^TM^ mice were injected subcutaneously at the right lower flank with MC38 or RM-1 cells (10^6^ cells suspended in 100 μL of PBS) as indicated for tumor development. Tumors were allowed to establish until tumor volume reached the indicated size. Mice were then randomized and dosing, as described for each experiment, was initiated on the same day as randomization. Tumor volumes and body weight were measured in a blinded fashion at least twice a week. Tumor volumes were calculated using the formula: V = (L × W × W)/2, where V is tumor volume, L is tumor length (the longest tumor dimension), and W is tumor width (the longest tumor dimension perpendicular to L). Individual animals were removed from the study as their tumor volumes measured greater than 3000 mm^3^. Where indicated mice with a complete response on day 35 post randomization were re-challenged on day 43. For the re-challenge experiment, MC38 tumor cells (10^6^ in 100 μL PBS) were implanted subcutaneously in the lower left (opposite) flank region. Tumor cells were also implanted in 6-8 week old naïve, WT C57BL/6 mice (Shanghai Lingchang Biotechnology Co., Ltd. (Shanghai, China)) as a positive control for tumor growth at the time of the re-challenge. Mice and any tumor progression were observed for another 30 days.

Single cell suspensions from tumors were prepared using the murine tumor dissociation kit (Miltenyi, Bergisch Gladbach, Germany) and GentleMACS^TM^ Octo Dissociator with Heaters set to the dissociation program (37_c_m_TDK_1). Single cell suspensions from lymph nodes were pushed through a 70 μm cell strainer using the plunger from a 5 mL syringe. Single cell suspensions were then incubated at room temperature with 1× RBC lysis buffer for 90 seconds. RBC lysis was then quenched, washed, strained through a 70 μm cell strainer, and counted. For cell staining cells were first blocked with 1 μg/ml Fc-Block (mouse Fc Block, BD Biosciences, San Jose, CA, USA) for 15 minutes at 4°C. Cells were then stained with surface antibodies for 30 minutes on ice in the dark. Cells were then washed twice and fixed with Fixation/Permeabilization working solution (eBioscience, San Diego, CA, USA) and then washed twice with 1× Permeabilization buffer (eBioscience, San Diego, CA, USA), after which intracellular staining was performed in 1× Permeabilization buffer. Following intracellular staining, cells were washed twice with 1× Permeabilization buffer and data was collected using a flow cytometer (LSRFortessa X-20, BD Biosciences, San Jose, CA, USA). Data were analyzed using Kaluza or FlowJo 10. Gating strategy is available in the supplemental information (**Fig. S7**).

Experiments using hCTLA-4/hPD-1 double knock-in HuGEMM^TM^ mice on the BALB/c background (GemPharmatech Co., Ltd; 62) were generated by crossing mice in which exon 2 of murine *Ctla4* was replaced with exon 2 from human *CTLA4* with mice in which exon 2 and 3 of murine *Pdcd1* were replaced with exon 2 and 3 of human *PDCD1*. The murine toxicity study using these mice was performed at Crown Bioscience (Taicang Jiangsu Province, China). Female mice were 4-5 weeks old at start of dosing and were treated with PBS, pembrolizumab (15 mg/kg), ipilimumab (10 mg/kg) plus pembrolizumab, or GIGA-564 (10 mg/kg) plus pembrolizumab every 3 days for 9 doses, and mice were euthanized 10 days after completion of dosing and tissues collected for histopathology analysis.

In graphs where data are shown longitudinally with lines connecting data points the measurements were taken repeatedly from the same group of animals over time.

### Epitope mapping

Identification of the key energetic residues involved in binding was performed by Integral Molecular (Philadelphia, PA, USA) (44). Full-length human CTLA-4 with a mutation to reduce internalization (Y201G) (63) was cloned into a transient expression vector, with each extracellular position (36–161) individually mutated to alanine (or alanine mutated to serine). The mutant library was arrayed in 384-well microplates and transiently transfected into HEK-293T cells. The optimal staining concentrations of ipilimumab, GIGA-564, and L3D10 control (BioLegend, San Diego, CA, USA) were determined using wild-type CTLA-4, then applied to the CTLA-4 mutant library. Antibody binding was detected using goat-anti-human IgG-Alexa fluor-488 or anti-human IgG F(ab’)2-Alexa fluor-488 (Jackson ImmunoResearch, West Grove, PA, USA), and mean cellular fluorescence was determined using flow cytometry (Intellicyt iQue, Ann Arbor, MI, USA). Mutated residues were considered critical if mutation resulted in significant loss of binding to the test antibody compared to the control antibody.

### *In vitro* Treg activation assay

Anti-CD3 (OKT3, Ultra-LEAF purified, BioLegend, San Diego, CA, USA) with and without CD80 (R&D Systems rhCD80-Fc, 10107-B1-100, Minneapolis, MN, USA) were covalently linked to the surface of M-450 Tosyl beads (Dynabead M-450, 140130, Thermo Fisher Scientific, Waltham, MA, USA) according to the manufacturer’s guidelines. To confirm that the surfaces of each bead type were properly coated, fluorescent antibodies were used to detect either the mouse Fc (anti-CD3) or the CD80 on the bead surface via flow cytometry.

Donor-matched human Treg and Tconv cells (Stemcell Technologies, 70096, Vancouver, BC, Canada) were thawed and stained with CellTrace Violet (C34571, Thermo Fisher Scientific, Waltham, MA, USA) according to the manufacturer’s protocol. 10^5^ cells were then cultured with 10^5^ beads as indicated in Complete Iscove’s Modified Dulbecco’s Medium (IMDM, Thermo Fisher Scientific, Waltham, MA, USA) with non-essential amino acids, sodium pyruvate, Glutamax, and 5% human antibody serum at 37°C, 5% CO_2_ in a 96-well U-bottom plate (Falcon-Corning U-bottom tissue culture plates, VWR, Radnor, PA, USA). In conditions where rhCD80-Fc (Abatacept, BMS, Princeton, NJ, USA) or aCTLA-4.28 were used, these were mixed with beads first prior to being added to the cells. Test samples with both rhCD80-Fc and aCTLA-4.28 were mixed and incubated together prior to mixing with beads and then cells. After a 4 day incubation, cells were washed, stained, and analyzed using the CytoFLEX LX (BD Biosciences, San Jose, CA, USA). Flow cytometry data were analyzed in FlowJo v10.7.1 (BD Biosciences, San Jose, CA, USA).

### FcR effector activity bioassays

Fc effector activity bioassays for mFcγRIIIa (CS1779B08), mFcγRIV (M1151), hFcγRIIb (CS1781E02), hFcγRIIa-H variant (G9981), hFcγRIIa-R variant (CS1781B08), hFcγRIIIa-V variant (G7011), and hFcγRIIIa-F variant (G9791) were purchased from Promega Corporation (Madison, WI, USA). The assays were carried out following the manufacturer’s instructions. Briefly, CHO target cells stably expressing human CTLA-4 with the Y201G mutation to enhance cell surface expression (63) were suspended in RPMI 1640 + 4% FBS media with either GIGA-564, ipilimumab, or GIGA-564 with the LALA-PG mutation to eliminate Fc function (64), and incubated at 37°C for 30 minutes. The following starting concentrations and dilution factors were used for each assay: 1 μg/mL, 2.5-fold dilution series (mFcγRIIIa, hFcγRIIIa-V variant), 500 μg/mL, 2.5-fold (mFcγRIV), 3 μg/mL, 3-fold (hFcγRIIb, hFcγRIIa-H and R variant), or 10 μg/mL, 2.5-fold (hFcγRIIIa-F variant). Jurkat/NFAT-Luc effector cells solely expressing one type of FcγR were added to each well (effector:target ratio was 5:1) and incubated at 37°C for 6 hours. Luciferase activity was measured by using the included Bio-Glo Luciferase Assay Reagent with the SpectraMax i3x or iD3 plate readers (Molecular Devices, San Jose, CA, USA). Luciferase activity measured in relative luminescence units (RLU) were plotted against the concentration of CTLA-4 mAbs. The IC50 value of each mAb was calculated by logistic regression using Prism (GraphPad, San Diego, CA, USA).

### Determination of pH-sensitivity of antibodies

96-well plates (Nunc, MaxiSorp, flat bottom, uncoated, BioLegend, San Diego, CA, USA, 423501) were coated with 0.5 μg/mL rhCTLA-4-Fc chimera (7268-CT, R&D Systems, Minneapolis, MN, USA) diluted in 1× PBS pH 7.0 and incubated at 2-8°C overnight. Plates were washed with 1× PBST and blocked with PBST + 1% BSA (PBSTB). Purified CTLA-4 mAbs were diluted in 10 mM sodium phosphate + 150 mM NaCl adjusted to pH 4.0, 5.0, 6.0, or 7.0 then added to the plate for 1 hour. Unbound antibodies were washed away with PBST and bound antibodies detected with 0.5 μg/mL HRP-conjugated goat anti-human constant kappa (2060-05, Southern Biotech, Birmingham, AL, USA) diluted in PBSTB. Plate was washed with PBST once more, then TMB substrate (1-Step™ Ultra TMB-ELISA Substrate Solution, Thermo Fisher Scientific, Waltham, MA, USA) was added and developed for approximately 1 minute before stopping the reaction with 1 N HCl. Absorbance at 450 nm was measured using a spectrophotometer (i3x, Molecular Devices, San Jose, CA, USA) and plotted using Prism (GraphPad, San Diego, CA, USA).

### Generation of cell pools stably expressing GIGA-564

GIGA-564 was expressed from a variant of the pCDNA5/FRT mammalian expression vector (Thermo Fisher Scientific; Waltham, MA, USA). The vector has a promoter for glutamine synthetase selection and uses an EF1α promoter to drive light chain expression, followed by a BGH polyA sequence and a CMV promoter to drive expression of the heavy chain, followed by a second BGH polyA sequence. The antibody expression construct was built using GeneBlocks (Integrated DNA Technologies, Coralville, IA, USA) and NEBuilder® HiFi DNA Assembly Master Mix (New England BioLabs, Ipswich, MA, USA). The construct was amplified in NEB 10-beta *E. coli* and purified with the ZymoPURE™ Plasmid Maxiprep Kit (Zymo Research, Irvine, CA, USA). It was then linearized for transfection by digesting 150 μg of DNA with 3,000 units PvuI-HF restriction enzyme (New England Biolabs, Ipswich, MA, USA), precipitated, and washed.

Sigma Aldrich’s CHOZN cells were cultured in EX-CELL CD CHO Fusion medium (MilliporeSigma, Burlington, MA, USA) supplemented with 6 mM GlutaMAX (Gibco, Waltham, MA, USA), shaking at 37°C, 5% CO_2_, 125 RPM (25 mm throw) for 7 days prior to electroporation. The day before electroporation, suspension cells were seeded at 0.5 × 10^6^ viable cells/mL (vc/mL). Linearized plasmid was transfected and pulsed using CM-150 on the Amaxa 4D Nucleofector (Lonza, Basel, Switzerland) and SE Cell Line 4D Nucleofector X kit (Lonza, Basel, Switzerland). Each cuvette had 4 μg of DNA and 10^7^ CHOZN cells concentrated in SE cell solution. Post-electroporation, the transfected cells were transferred to T-75 flasks at a 2 × 10^6^ vc/mL density for a 1-day recovery in EX-CELL CD CHO Fusion medium (MilliporeSigma, Burlington, MA, USA) with 6mM GlutaMAX (Gibco; Waltham, MA, USA). After 1-day recovery, transfected cells were pelleted and resuspended into 80% EX-CELL CD CHO Cloning medium (MilliporeSigma, Burlington, MA, USA) and 20% EX-CELL CD CHO Fusion medium (MilliporeSigma, Burlington, MA, USA) and plated into 96-well plates (non-TC treated, flat bottom, Greiner One-Bio, Kremsmünster, Austria) at 5,000 cell per well and incubated 37°C, 5% CO_2_ with humidity.

After 4-5 weeks, confluent transfected wells (minipools) were screened for antibody concentration using the Gator bio-layer interferometry system and Protein-A biosensors (GatorBio, Palo Alto, CA, USA). Minipools with the highest antibody titer were scaled-up using EX-CELL CD CHO Fusion medium in static 24-well TC-treated plates incubated at 37°C, 5% CO_2_ with humidity. Minipools continued to expand and shaker adapted to shaking 6-well plates (non-TC treated, flat bottom, Greiner One-Bio, Kremsmünster, Austria) on a 19 mm shaking platform set at 145 RPM, 37°C, 5% CO_2_, with humidity. Material from an 8-day terminal batch in 24-well plates (TC treated, flat bottom, Corning, Corning, NY, USA) was screened using the Gator system as a second method to screen minipools, in order to identify the high producers. Cultures were expanded into shaking 50 mL conical tubes (Midsci, St. Louis, MO, USA) incubated at 37°C, 5% CO_2_, 80% humidity, shaking at 225 RPM with 25 mm throw. Top minipools were combined to generate enriched pools (EPs).

EPs were cultured in shake flasks at 125 RPM, 25 mm throw, seeded at 0.2-0.4 × 10^6^ viable cell density (VCD)/mL for 3-4 days. EPs were then inoculated in EX-CELL Advanced Fed Batch medium at 0.4 × 10^6^ VCD/mL in 1 L shake flasks (Corning, Corning, NY, USA) with a vent cap and working volume of 200 mL. Incubator conditions were held constant throughout fed-batch at 37°C, 5% CO_2_, 80% humidity, and shaking at 125 RPM (25 mm throw). Beginning day 3 of fed-batch to harvest, growth, viability, and metabolite concentrations were measured offline. Glucose was supplemented when concentration dropped below 4 g/L, up to 6 g/L using 45% glucose stock (Corning, Corning, NY, USA). Cultures were fed days 3, 5, 7, 9, and 11 with EX-CELL Advanced CHO Feed 1 (MilliporeSigma, Burlington, MA, USA) (4% of working culture volume) and CellVento 4Feed COMP (MilliporeSigma, Burlington, MA, USA) (2% of working culture volume). Cultures were harvested when viability dropped below 80% and cell culture supernatant was clarified by centrifugation and filtered. Protein was purified using Protein A eluted with 100 mM acetate pH 3. One EP was used for subsequent analysis (EP-1).

### Glycan analysis

This analysis was performed by the CRO Bionova Scientific (Fremont, CA, USA). 150 μg of antibody was digested with PNGase F (Prozyme, Hayward, CA, USA) to liberate glycans. Glycans were isolated and labeled with a fluorescent molecule using a GlykoPrep InstantAB labeling kit (Prozyme, Hayward, CA, USA) following the manufacturer’s instructions. Labeled glycans were injected over a 3.5 μm, 2.1 × 150 mm XBridge Amide column (Waters, Milford, MA, USA) on a UPLC (Dionex Ultimate 3000, Waters, Milford, MA, USA) with a fluorescent detector. Peaks were identified based on known glycans from the Glyko InstantAB biantennary and high-mannose partitioned library standards (Prozyme, Hayward, CA, USA), with the total percent of each glycoform reported as the integrated area under the identified peak divided by the sum of the AUC (area under the curve) for all peaks.

### Skin inflammation, colonic epithelial and heart toxicity scoring

Hematoxylin and eosin (H&E)-stained sections of skin from near the whisker region was scored as follows 1: 1-3 small foci of lymphocyte aggregates per section, 2: 4-10 small foci or 1– 3 intermediate foci, 3: 4 or more intermediate or the presence of large foci, 4: marked interstitial fibrosis in parenchyma and large foci of lymphocyte aggregates.

The colonic epithelial score was determined by assessment of H&E-stained colon rolls. To determine this score, after reviewing the whole slide the four regions in which damage was most severe were selected for scoring, and the score from all four areas was added together to determine the cumulative score. To determine the score for individual areas, both the epithelial structures affected and the consistency of the damage were taken into account. For this the damage to epithelial structures was scored as 0: no lesions, 1: mucosa damage, 2: submucosa damage, 3: muscularis/serosa damage. The consistency of damage in each area was scored as 1: focal, 2: patchy, 3: diffuse. Both scores were then multiplied together to determine the area score.

For the heart pathology score, lymphocyte infiltration on H&E-stained heart sections in pericardium, right or left atrium, base of aorta, and left or right ventricle each count for 1 point each. The number of CD45^+^ cells were counted on heart formalin-fixed paraffin-embedded (FFPE) sections stained with CD45. The visual fields for inflammatory cell infiltration analysis were randomly selected, and the average score of three fields for each sample was determined.

### Kidney pathology staining and scoring

Tissues were collected and were placed in 10% buffered neutral formalin and then paraffin embedded. Sectioning, staining, and scoring was performed by Allele (San Diego, CA, USA). Briefly, blocks were cut in 5 μm sections that were placed on glass slides for anti-IgG (UltraPolymer Goat anti-Mouse heavy and light chain IgG-HRP, Cell IDx, San Diego, CA, USA) or anti-C3 (EPR19394, Abcam, Cambridge, UK) immunohistochemistry (IHC) staining. Stained slides were prepared as digital images.

A board-certified veterinary pathologist with experience in laboratory animals and toxicologic pathology, who was blinded to the study, evaluated the anti-IgG slides for location, intensity, and percent of positive staining. Findings were scored on a scale of 1 to 4 for intensity (0: negative, 1: minimal or slightly positive, 4: very dark), and as a percent of the positive cells in the glomeruli (after reviewing at least 5 glomeruli).

### Statistical analysis

Comparison of changes in tumor volume measured longitudinally in multiple groups was determined using a linear mixed effects model with treatment group and day as fixed effects and animal identifier as a random effect, to account for the dependence of repeated measures. All tests were two-sided without adjustments to type I error rates. These analyses were conducted using R version 3.6.2 (65).

For the re-challenge study, tumor volume (in mm^3^) was analyzed using a linear mixed effects model including treatment group and day as fixed effects and animal identifier as a random effect, to account for repeated measures. Statistical comparisons were made using the Wald test against the isotype control group for the initial challenge and against the naïve control group for the re-challenge portion of the study. All tests were two-sided with an alpha level of 0.05.

Adjusted p-values for comparison of skin inflammation, colonic epithelial damage, CD45^+^ cell infiltration into the heart, heart pathology score, colon length, spleen weight, and kidney Ig or C3 deposition were calculated using the Dunn’s test and the Benjamini-Hochberg step-down procedure to account for multiple comparisons (66). Analyses were performed using R version 3.6.2 (65). To determine if there was any statistical difference in the percent body weight induced by pembrolizumab or pembrolizumab plus a CTLA-4 mAb in comparison to control, a linear mixed effects model was fit on change in body weight including day and treatment group as fixed effects and animal identifier as a random effect, to account for repeated measurements within animal.

For flow cytometry data, the Wilcoxon rank sum test was used to determine statistical significance, with an alpha level of 0.05, two-sided.

## ACKNOWLEDGEMENTS

The authors thank Penny Chen, Tina Zhang, Davy Ouyang, Mingfa Zhang, Michael Boice, Mary Topalovski, and Katharina Hodges (Crown Bioscience) for assistance with *in vivo* studies. The authors also thank Abbas Hussain and Anthony Wong (Allele Biotechnology) and Likun Zhang (Crown Biosciences) for assistance with IHC.

## SUPPLEMENTARY MATERIALS

Fig. S1. ***In vitro* characterization of scFvs reformatted as full-length antibodies.**

Fig. S2. **Validation of N297Q mutants binding to cell-surface CTLA-4.**

Fig. S3. **Co-stimulation enhances Treg proliferation.**

Fig. S4. **Anti-CTLA-4s deplete intratumoral Tregs in hCTLA-4 KI mice.**

Fig. S5. **GIGA-564 induces more FcR signaling than ipilimumab.**

Fig. S6. **GIGA-564 results in less toxicity than ipilimumab in murine models.**

Fig S7. **Gating strategy for analysis of T cell populations in hCTLA-4 KI mice bearing MC38 tumors and treated with anti-CTLA-4s.**

Fig S8 - 10. Additional gating strategy figures.

Table S1. **In vitro characterization of scFvs reformatted as full-length antibodies.**

Table S2. **GIGA-564 has minimal ability to block CD80 or CD86 from binding CTLA-4.**

Table S3. **Fc receptor signaling of GIGA-564 versus ipilimumab.**

Table S4. **CTLA-4 antibody sequences.**

## AUTHOR CONTRIBUTIONS

Conceptualization, E.L.S., A.S.A., and D.S.J.; methodology, E.L.S., K.P.C., E.K.W., M.A.A., A.R.N., S.K.S., J.F.S., L.T., N.P.W., and Y.W.L.; software, Y.W.L.; investigation, E.L.S., K.P.C., E.K.W., M.A.A., E.B., Y.Y.C., G.L.C., C.E., B.K.G., A.G., J.L., R.L., V.A.M., R.A.M., A.R.N., J.S., S.K.S., J.F.S., K.S., B.T., L.T., N.P.W., and Y.W.L.; data curation, E.L.S., K.P.C., E.K.W., M.A.A., A.R.N., L.T., Y.W.L., and A.S.A.; writing—original draft preparation, E.L.S. and D.S.J.; writing—review and editing, E.L.S., A.S.A., and D.S.J.; visualization, E.L.S., K.P.C. and Y.W.L.; supervision, E.L.S., K.P.C., A.S.A., and D.S.J.; project administration, E.L.S., A.S.A., and D.S.J.; funding acquisition, D.S.J.

## DATA AND MATERIALS AVAILABILITY

All data associated with this study are available in the main text, as supplementary materials, or upon reasonable request (Figure 1A, 2D, 4, 7B-C, S1A, C, S2 - S4, S6C-E, and Table S1-3. Certain materials, such as proprietary antibodies, antibody expression constructs, and cell lines, can be made available for non-commercial research, subject to a material transfer agreement.

