## Supplemental Information for "Lack of blocking activity in anti-CTLA-4 antibodies reduces toxicity, but not anti-tumor efficacy"

### SUPPLEMENTAL FIGURES

#### Supplemental Figure 1

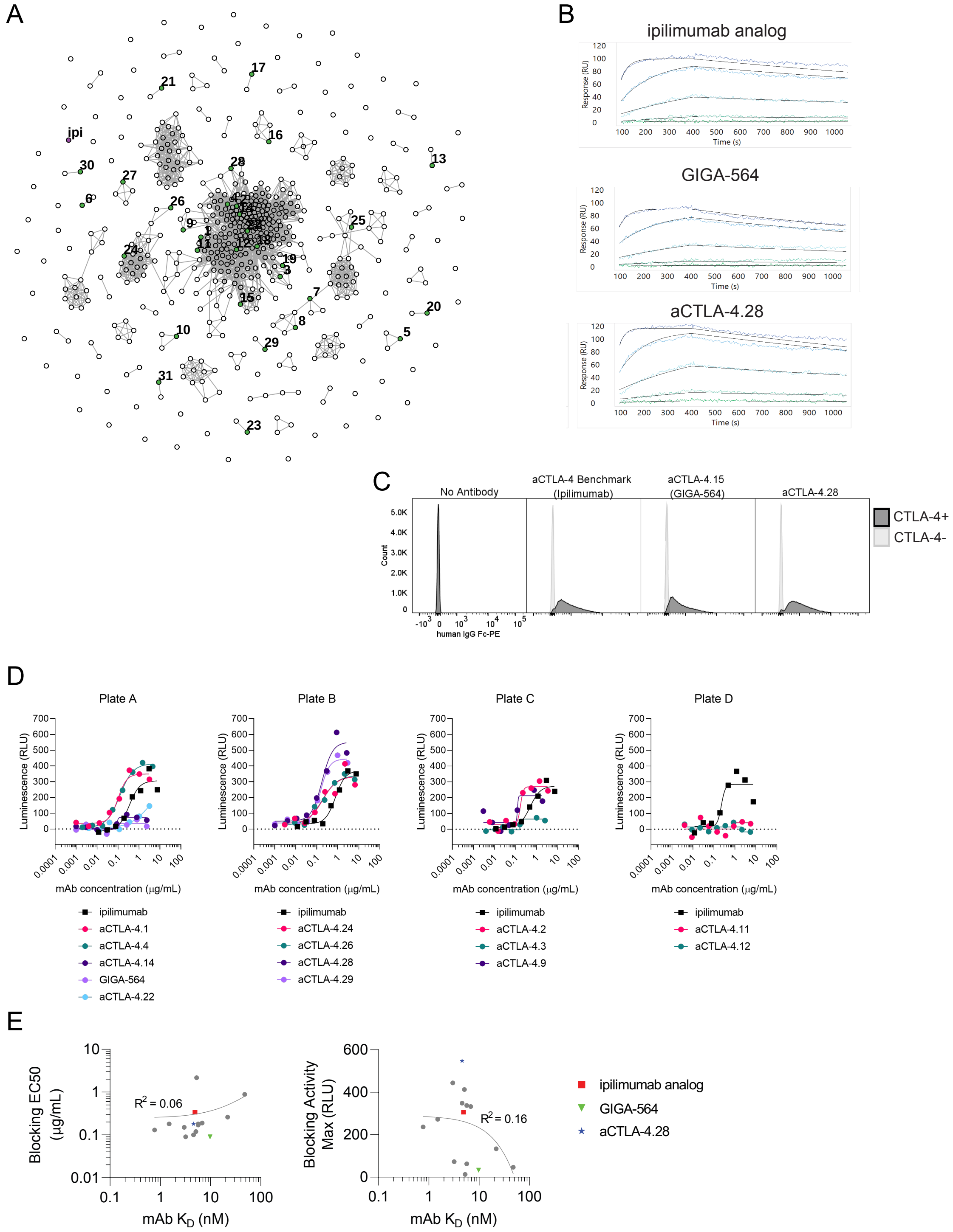

**Fig. S1. *In vitro* characterization of scFvs reformatted as full-length antibodies.**

**A**. Clonal cluster analysis for the FACS-enriched anti-CTLA-4 scFv clones. Each node represents an scFv clone (full-length IgK+IgH). We computed the total number of amino acid differences between each pairwise alignment of scFv sequences. Edges indicate pairwise alignments with ≤9 amino acid differences (Clustergram modified from Fig. 7 of Asensio et al., 2019). The scFv sequence for ipilimumab (purple, ipi) was included for comparison. scFv clones for full-length antibodies described in this manuscript are in green and labeled with ID numbers. **B**. The affinity of the indicated antibody to soluble CTLA-4 was determined by SPR (Carterra). Plots show association and dissociation signals with a 5-fold dilution series of antigen starting at 500 nM. **C**. A 50:50 mixture of CTLA-4$^{+}$ and CD27$^{+}$ (CTLA-4$^{-}$) CHO cells were stained with 10 μg/ml of the indicated antibody. An anti-human IgG secondary antibody conjugated to PE was used to detect cells labeled with the indicated anti-CTLA-4 antibody, while anti-CD27-FITC identified CD27$^{+}$ CHO cells. Histograms show staining of CTLA-4$^{+}$ (CD27-FITC$^{-}$) or CTLA-4$^{-}$ (CD27-FITC$^{+}$) cells by the indicated antibody as determine by flow cytometry. See **Fig S8** for gating strategy. **D**. The cell-based CTLA-4 Blockade Bioassay (Promega) involved co-culturing CTLA-4-expressing Jurkat cells with Raji cells, which naturally express CD80 and CD86, in the presence of the indicated mAbs. mAbs that bind to CTLA-4 and block the ability of CTLA-4 to interact with CD80/CD86 lead to CD28 pathway-activated luciferase expression. Plot shows the amount of luciferase expression induced (relative luciferase units; RLU) when cells were cultured with a titration series of the indicated antibody. Due to constraints on sample size in each Promega bioassay kit, this set of aCTLA-4 mAbs was analyzed using multiple plates. To control for plate-to-plate variability in maximum signal, the ipilimumab analog was run on each plate, and a representative sample was used to calculate the EC50 and maximum signal for ipilimumab in **Table S1**. The plate in which each antibody was tested is indicated in **Table S1**. Plate A and B were run at the same time, while Plate C and Plate D were run separately at later times. Each point represents a single measurement. **E.** Correlation between antibody affinity and cell-based assay activity. Antibody affinity (KD) for CTLA-4 was compared to the blocking EC50 and the RLU maximum signal; no correlation was found for either when the data was fit by linear regression. Each data point represents a single antibody, and the following antibodies are distinguished as follows: ipilimumab analog (red square), GIGA-564 (green inverted triangle), and aCTLA-4.28 (blue star). Data is from (**D**) and **Table S1**.

Supplemental Figure 2

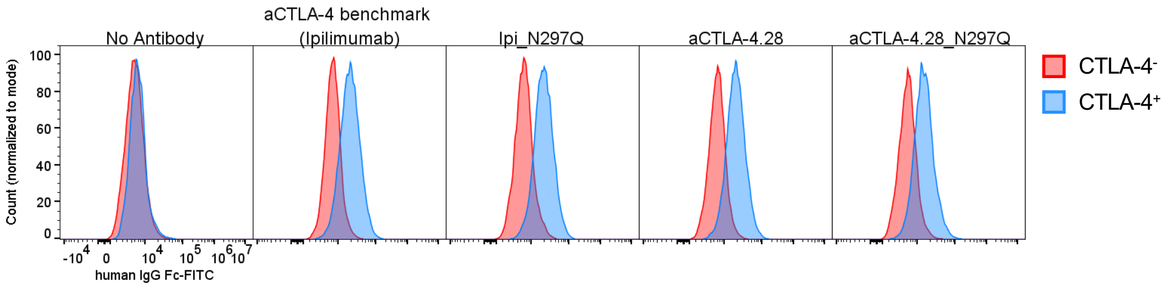


**Fig. S2. Validation of N297Q mutants binding to cell-surface CTLA-4.**

CHO cells with and without human CTLA-4 expression were incubated with 10 µg/mL of the indicated antibody, then an anti-human IgG secondary antibody conjugated to FITC was used to detect cells bound by the indicated CTLA-4 antibody. Histograms show staining of CTLA-$4^{+}$ or CTLA-$4^{-}$ cells by the indicated antibody as determine by flow cytometry. Gating strategy is available in **Fig. S9**.

Supplemental Figure 3

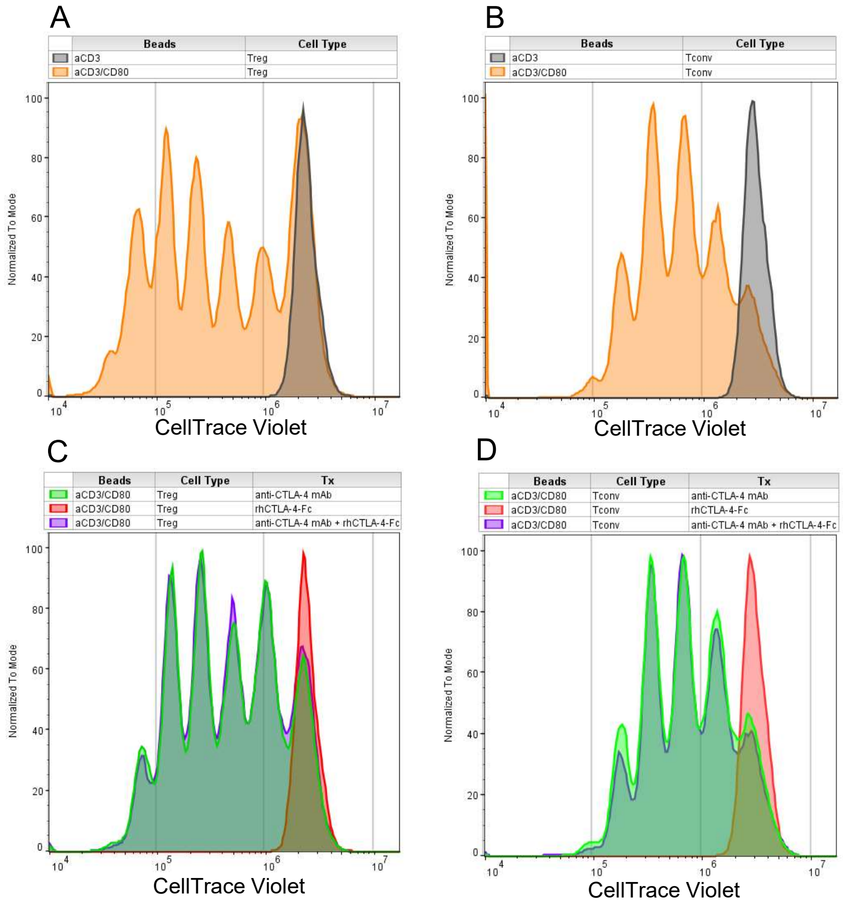


**Fig. S3. Co-stimulation enhances Treg proliferation.**

Histograms show CellTrace Violet signal (gated on Live, $CD3^{+}CD4^{+}$ cells) of **A.** Treg or **B.** Tconv cells cultured in the presence of M-450 Tosyl activated beads coated with anti-CD3 antibody with and without CD80, or **C.** Treg or **D.** Tconv cells activated with M-450 Tosyl activated beads coated with anti-CD3 antibody plus CD80 in the presence of rhCTLA-4 (Abatacept) and/or anti-CTLA-4 mAb (aCTLA-4.28). Data is from a single donor in a single experiment. No technical replicates were performed due to the limited number of available cells.

Supplementary Figure 4

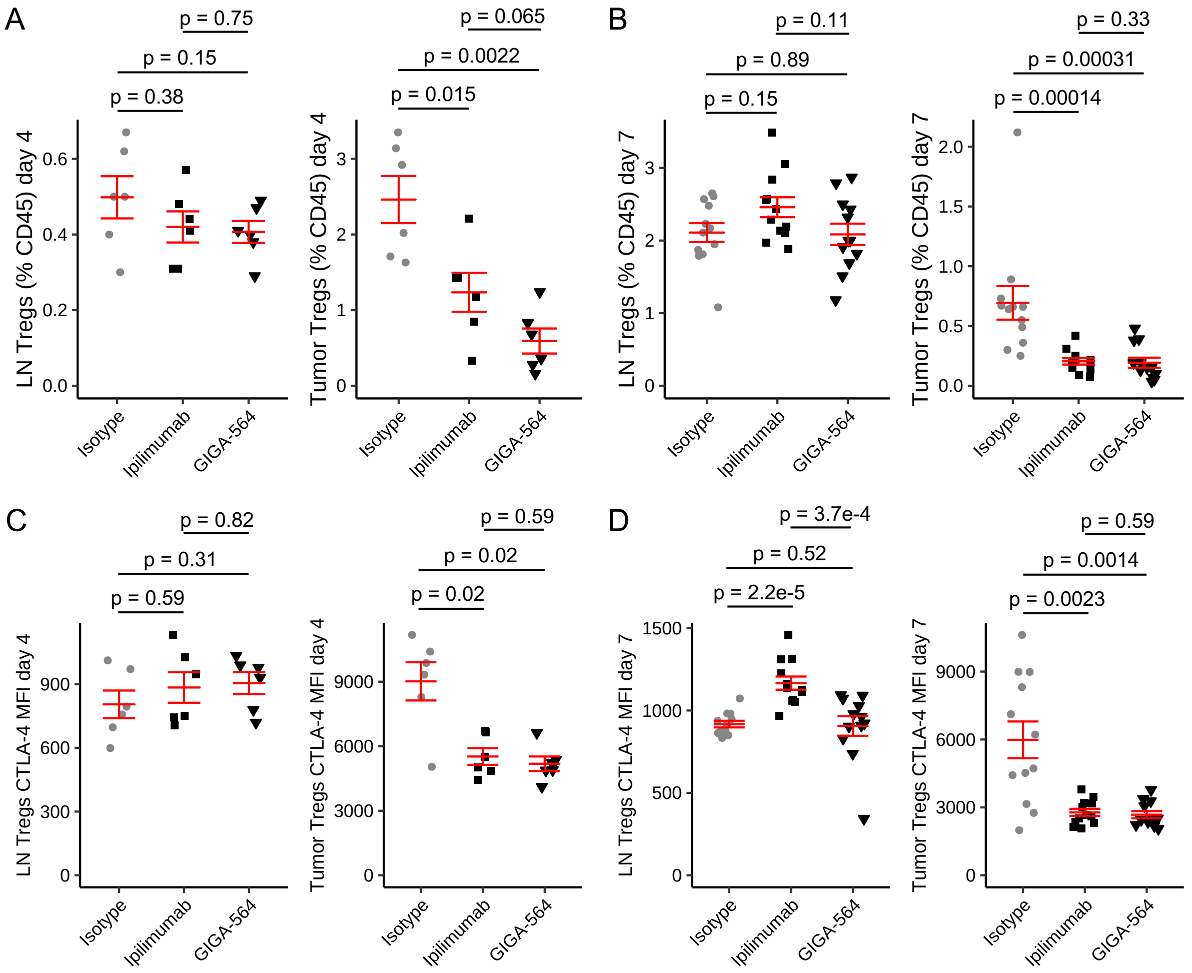


**Fig. S4. Anti-CTLA-4s deplete intratumoral Tregs in hCTLA-4 KI mice.**

Flow cytometry analysis of cells from MC38 tumor bearing hCTLA-4 KI mice receiving CTLA-4 mAbs. **A** and **C.** 6 mice per group were treated with 5 mg/kg hIgG1 isotype control, ipilimumab, or GIGA-564 on day 0 and 3, and cells were analyzed on day 4. **B** and **D.** 12 mice per group were treated with 5 mg/kg hIgG1 isotype control, ipilimumab, or GIGA-564 on day 0, 3, and 6, and cells were analyzed on day 7. Two lymph node samples in the GIGA-564 group were excluded from analysis due to low cell count. **A-B.** Frequency of Tregs (Live, $CD45^{+}TCR\beta^{+}CD4^{+}FOXP3^{+}$, as percentage of $CD45^{+}$ cells) in lymph nodes (left) and tumor (right) in each treatment group. **C-D.** Geometric mean fluorescence intensity (MFI) of intracellular CTLA-4 in Tregs in LN (left) and tumor (right). P-values were calculated using a two-sided Wilcoxon rank sum test. Red lines depict mean +/- SEM.

#### Supplemental Figure 5

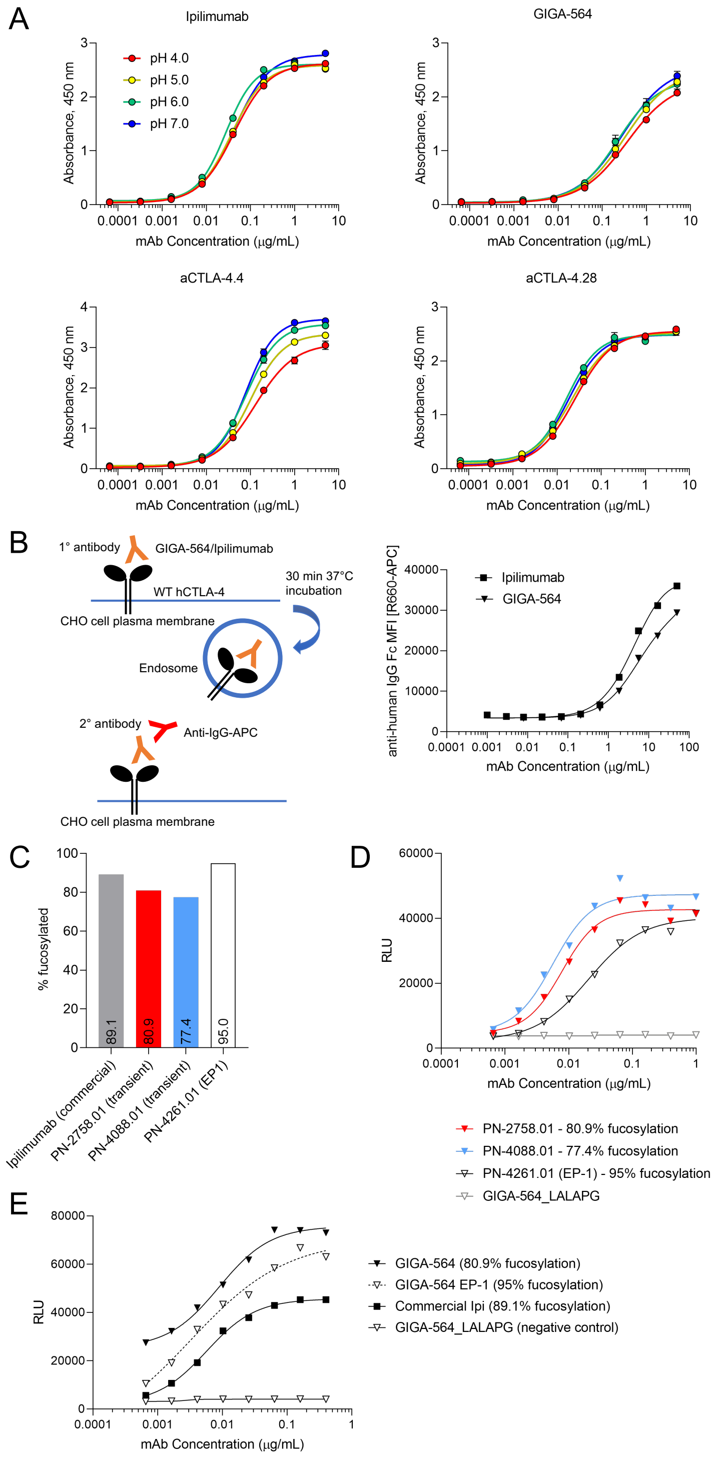


**Fig. S5. GIGA-564 induces more FcR signaling than ipilimumab.**

**A.** Purified CTLA-4 antibodies were diluted to a starting concentration of 5 μg/mL for an 8-point, 5-fold titration series in different pH buffers, then added to wells coated with rhCTLA-4-Fc. Bound antibodies were detected with anti-constant kappa-HRP and measured for absorbance

at 450 nm. Each point is the average of two technical replicates. **B.** CHO cells expressing wildtype hCTLA-4 incubated with titrations of ipilimumab or GIGA-564 at 37°C to allow for internalization. The amount of antibody remaining on the surface was then determined by staining with anti-human IgG Fc. Each point is the median fluorescent intensity (MFI) of the anti-human IgG Fc-APC signal. Gating strategy is available in **Fig. S10**.
**C-D**. GIGA-564 from three different productions was tested for fucosylation levels (bar graphs each depict a single data point) (**C**) and human FcγRIIIA signaling (**D**). PN-2758.01 and PN-4088.01 were generated from transient transfection of ExpiCHO cells, while PN-4261.01 was from a stably-expressing pool of CHOZN clones (EP-1, enriched pool 1). **C.** The fucosylation level, as determined by UPLC analysis, is shown for each sample. **D-E**. Graph shows human FcγRIIIA (V variant) signaling as determined in a cell-based assay for the indicated three preparations of GIGA-564 (**D**) or GIGA-564 and ipilimumab with the indicated amount of fucosylation (**E**). A negative control protein with mutations to disrupt Fc receptor binding (GIGA-564_LALAPG) was also tested in the reporter bioassays. Data shown are RLU emitting from effectors cells. Each point represents a single measurement.

Supplemental Figure 6

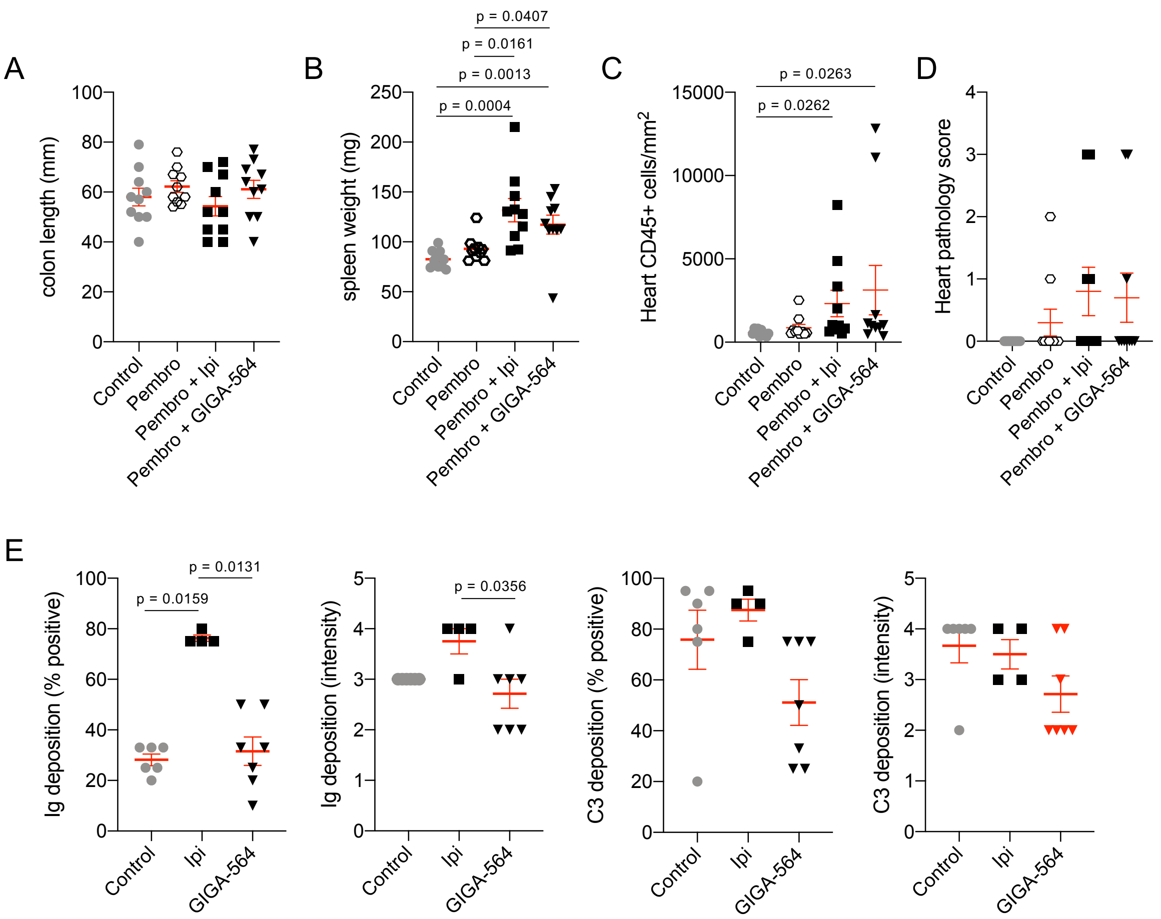


**Fig. S6**. **GIGA-564 results in less toxicity than ipilimumab in murine models.**

**A-D**. hCTLA-4/hPD-1 double KI mice on the BALB/c background between 4 to 5 weeks of age were treated with vehicle, pembrolizumab, pembrolizumab plus ipilimumab, or pembrolizumab plus GIGA-564 every 3 days for 9 doses. One week after the last dose mice were euthanized and tissues collected for pathology analysis. Graphs show colon length (**A**) or spleen weight (**B**) from mice of each group at time of euthanasia. The number of CD45$^{+}$ cells/mm$^{2}$ counted on randomly selected CD45 stained heart FFPE sections (**C**) or heart pathology score (**D**) as determined by a pathologist are shown. **E.** hCTLA-4 KI mice bearing MC38 tumors were treated with vehicle (PBS) or 5 mg/kg ipilimumab or GIGA-564 twice a week for 5 doses (**Fig. 3a**). On day 20 post initiation of treatment, mice were euthanized and kidneys were processed into formalin fixed paraffin embedded blocks (FFPE). FFPE sections were stained for anti-mouse immunoglobin or C3 and scored by a board-certified veterinary pathologist blinded from the study. Positive staining in glomeruli was along the capillary basement membranes. Graphs show percent of glomeruli positive for anti-murine IgG or C3 and the relative intensity of positive glomeruli; at least 5 glomeruli were examined in each section on 40x/high power. Red lines indicate means +/-

SEM. Adjusted p-values were calculated using the two-sided Dunn’s test applying the Benjamini-Hochberg step-down procedure to control for multiple comparisons.

Supplemental Figure 7A Tumor

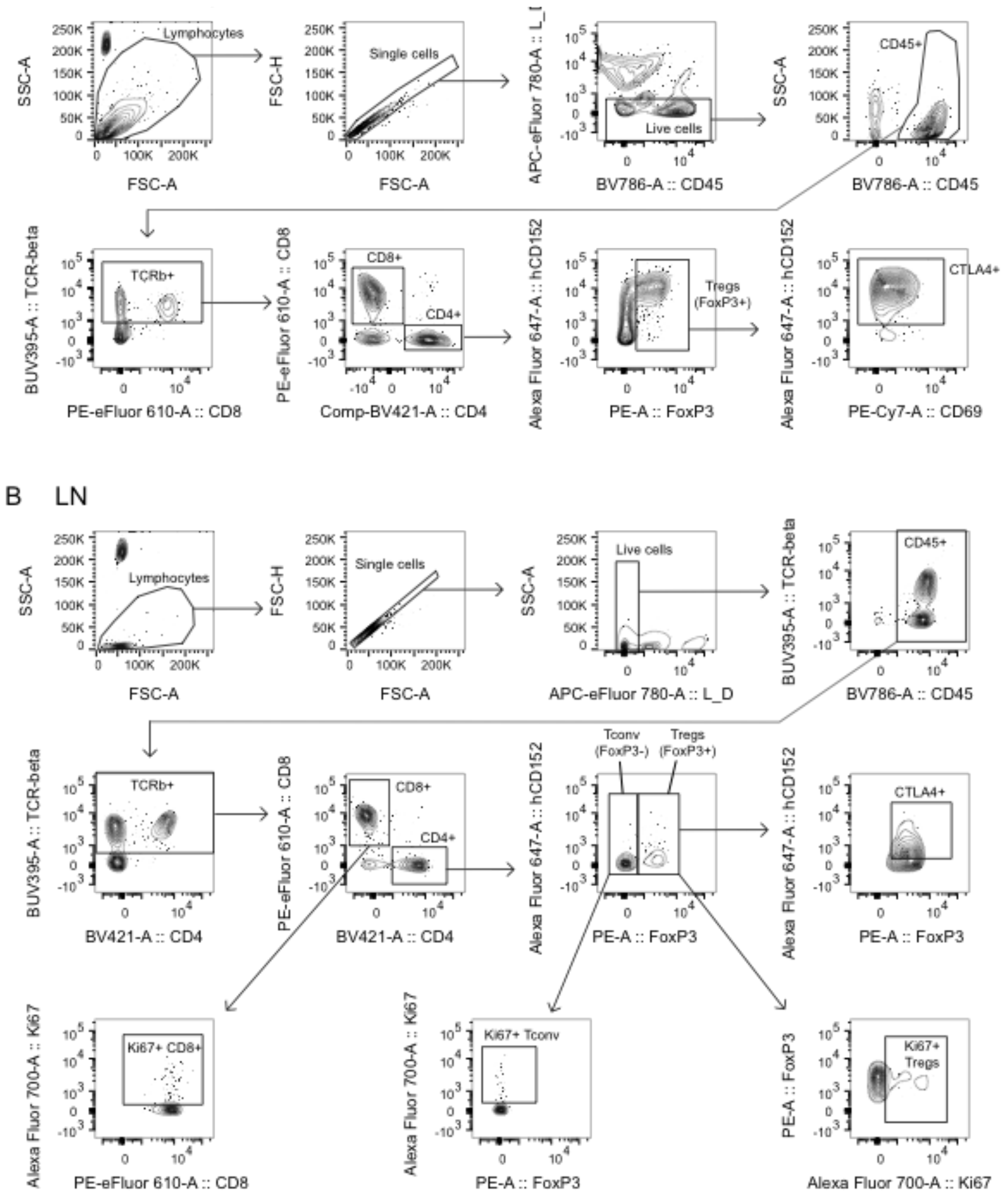

### Supplemental Figure 8 (Fig. S1C gating)

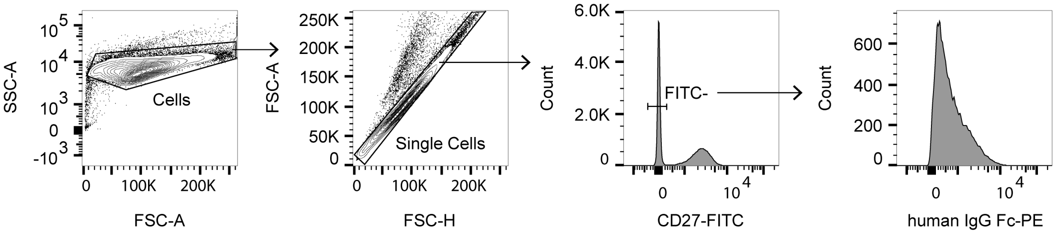


Samples were gated for cells by forward scatter and side scatter, then for single cells by forward scatter (height and area). Cells were then gated based on expression of CD27, where CD27+ cells are CTLA-4- and CD27- cells are CTLA-4+. The staining intensity of human IgG Fc-PE (which was used to detect binding of anti-CTLA-4 mAbs) on cells with (CD27-FITC-) and without (CD27-FITC+) CTLA-4 was then visualized using histograms.

### Supplemental Figure 9 (Fig. S2A gating)

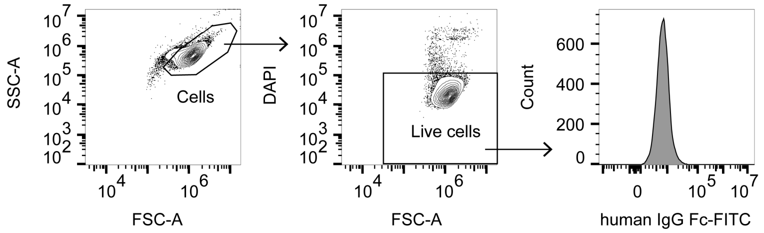


Samples were gated for cells by forward scatter and side scatter, then for live cells by low DAPI signal and forward scatter. Samples were then analyzed for staining intensity of human IgG Fc-FTIC for each antibody binding to cells with or without CTLA-4 expression.

### Supplemental Figure 10 (Fig. S5B gating)

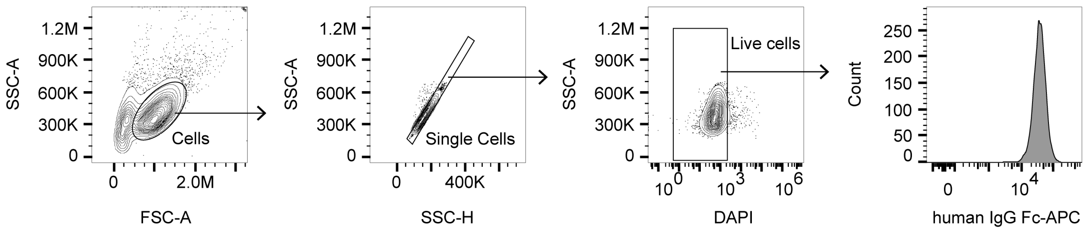


Samples were gated for cells by forward scatter and side scatter, then for single cells by side scatter (height and area), then for live cells by low DAPI signal and side scatter. Samples were then analyzed for staining intensity of human IgG Fc-APC for each antibody binding to cells.

1

### SUPPLEMENTAL TABLES

| Antibody | kon human CTLA-4 (M-1 s-1) | koff human CTLA-4 (s-1) | KD human CTLA-4 (nM) | MFI of CTLA-4+ CHO cells | MFI of CD27+ CHO cells | Cell-based blocking activity EC50 (µg/mL) | Cell-based blocking activity max signal (RLU) | Cell-based assay plate |
|---|---|---|---|---|---|---|---|---|
| Ipilimumab analog | 6.50E+04 | 3.20E-04 | 4.9 | 860 | 26.6 | 0.34 | 307 | A |
| aCTLA-4.1 | 8.50E+04 | 3.90E-04 | 4.6 | 1008 | 18.6 | 0.10 | 349 | A |
| aCTLA-4.2 | 1.90E+05 | 2.80E-04 | 1.5 | 1050 | 18.2 | 0.18 | 273 | C |
| aCTLA-4.3 | 6.40E+04 | 3.60E-04 | 5.7 | 854 | 15.3 | 0.17 | 63 | C |
| aCTLA-4.4 | 1.00E+05 | 5.20E-04 | 5.1 | 1123 | 11.7 | 0.12 | 413 | A |
| aCTLA-4.9 | 2.10E+05 | 1.60E-04 | 0.77 | 1328 | 18.5 | 0.13 | 237 | C |
| aCTLA-4.11 | 4.20E+03 | 2.00E-04 | 48 | 219 | 14.8 | 0.89 | 47 | D |
| aCTLA-4.12 | 5.10E+04 | 2.70E-04 | 5.3 | 1013 | 17.1 | 2.17 | 13 | D |
| aCTLA-4.14 | 1.30E+05 | 4.20E-04 | 3.2 | 1024 | 17 | 0.09 | 73 | A |
| aCTLA-4.15 (GIGA-564) | 5.40E+04 | 5.20E-04 | 9.8 | 717 | 15.9 | 0.09 | 33 | A |
| aCTLA-4.22 | 2.30E+04 | 5.10E-04 | 22 | 641 | 15.2 | 0.26 | 134 | A |
| aCTLA-4.24 | 6.30E+04 | 4.30E-04 | 6.8 | 934 | 26 | 0.19 | 333 | B |
| aCTLA-4.26 | 6.10E+04 | 3.50E-04 | 5.7 | 939 | 17.8 | 0.18 | 338 | B |
| aCTLA-4.28 | 9.00E+04 | 4.20E-04 | 4.6 | 1033 | 17.2 | 0.18 | 548 | B |
| aCTLA-4.29 | 7.60E+04 | 2.30E-04 | 3 | 999 | 16.3 | 0.15 | 444 | B |

**Table S1. *In vitro* characterization of scFvs reformatted as full-length antibodies.**

kon, koff, and KD of the indicated CTLA-4 mAbs for soluble human CTLA-4 was determined by SPR (Carterra). Geometric MFI of the indicated CTLA-4 mAbs bound to CTLA-4$^+$ or CD27$^+$ CHO cells as detected by a secondary human IgG antibody and flow cytometry. The ability of each aCTLA-4 mAb to block CTLA-4 binding to CD80/CD86 was determined by a cell-based assay (Promega). Due to constraints on sample size in each Promega bioassay kit, this set of aCTLA-4 mAbs was analyzed using multiple plates (plate letters correspond to Plate A-D in **Fig. S1D**). To control for plate-to-plate variability in maximum signal, the ipilimumab analog was run on each plate; a representative sample was used for this table.

| | CD80 | | | CD86 | | |
|---|---|---|---|---|---|---|
| Antibody | Max | Min | IC50 (µg/mL) | Max | Min | IC50 (µg/mL) |
| Ipilimumab | 0.96 | 0.04 | 0.07 | 1.04 | 0.09 | 0.07 |
| GIGA-564 | 0.97 | 0.18 | 2.23 | 1.07 | 0.36 | 0.30 |
| aCTLA-4.28 | 1.09 | 0.04 | 0.12 | 1.05 | 0.12 | 0.12 |

Max: maximum absorbance at 450 nm, normalized to prembrolizumab
Min: minimum absorbance at 450 nm, normalized to pembrolizumab

**Table S2. GIGA-564 has minimal ability to block CD80 or CD86 from binding CTLA-4.**

Summary of blocking ELISA data from **Fig. 2B**. Absorbance values from the detected CD80 or CD86 were normalized to signal from ELISA plates incubated with pembrolizumab. The data were then analyzed and fit by nonlinear regression. The normalized maximum and minimum signals and the IC50 for blocking are provided.

| FcR assay | Maximum luminescence (RLU) | | | EC50 (µg/mL) | |
|---|---|---|---|---|---|
| | Ipilimumab | GIGA-564 | GIGA-564_LALAPG | Ipilimumab | GIGA-564 |
| mFcγRIV | 990 | 2,557 | 297 | 2.21 | not determined* |
| mFcγRIII | 17,496 | 14,311 | 1,163 | 0.22 | 0.15 |
| hFcγRIIIa-V | 59,205 | 87,184 | 4,255 | 0.04 | 0.02 |
| hFcγRIIIa-F | 6,880 | 15,818 | 2,434 | 0.09 | 0.04 |
| hFcγRIIa-H | 973 | 802 | 210 | 0.05 | 0.04 |
| hFcγRIIa-R | 2,641 | 2,560 | 1,572 | 0.21 | 0.08 |
| hFcγRIIb | 5,801 | 5,173 | 978 | 0.17 | 0.11 |

*GIGA-564 EC50 not determined due to poor curve fit

**Table S3. Fc receptor signaling of GIGA-564 and ipilimumab.**

Summary of FcR Bioassay results from **Fig. 5**. Luminescence (RLU) data from the Bioassay were fit by nonlinear regression. The maximum signal and EC50 for FcR signaling are provided.

| Antibody | IgK variable sequence | IgG variable region sequence |
|---|---|---|
| aCTLA-4.1 | EIVLTQSPGTLSLSPGERATLSCRASQSVSYLAWYQQKPGQAPRLLIYGASSRATG<br>IPDRFSGSGSGTDFTLTISRLEPEDFAVYYCQQYGSSPWTFGQGTKVEIK | QVQLVESGGGVVQPGRSLRLSCAASGFTFSRYGMHWVRQAPGKGLEWVAVIWYDGRNKYYADSVK<br>GRFTISRDNSKNTLYLQMNSLRAEDTALYSCARAGELGPFDYWGQGTLVTVSSA |
| aCTLA-4.2 | EIVLTQSPGTLSLSPGERATLSCRASQSVSSSYLAWYQQKPGQAPRLLIYGASSRA<br>TGIPDRFSGSGSGTDFTLTISRLEPEDFAVYYCQQYGSSPFTFGPGTKVDIK | QVQLVESGGGVVQPGRSLRLSCIASGFTFSSYGMHWVRQAPGKGLEWVAVNWYDGSNKHYADSVK<br>GRFTISRDNSKNTLYLQMNSLRAEDTAVYYCARGGVWGPYFDYWGQGTLVTVSSA |
| aCTLA-4.3 | EIVLTQSPGTLSLSPGERATLSCRASQSVSSSYLAWYQQKPGQAPRLLIYGASSRA<br>TGIPDRFSGSGSGTDFTLTISRLEPEDFAVYYCQQYGSSPFTFGPGTKVDIK | QVQLVESGGGVVQPGRSLRLSCAASGFTLSSFGMHWVRQAPGKGLEWVAVIWYDGSNKYYADSVK<br>GRFIISRDNSKTALYLQMNSLRAEDTAVYYCARAHYFGAFDIWGQGTMVTVSSA |
| aCTLA-4.4 | EIVLTQSPGTLSLSPGERATLSCRASQSVSSSYLAWYQQKPGQAPRLLIYGASSRA<br>TGIPDRFSGSGSGTDFTLTISRLEPEDFAVYYCQQYGRSPFTFGPGTKVDIK | QVQLVESGGGVVQPGRSLRLSCAASGFTFSNYGMNWVRQAPGKGLEWVAVIWYDGRNKHYADSVK<br>GRFTISRDNSKNTLYLQMNSLRAEDTAVYYCARGGDWGPYFDYWGQGTLVTVSSA |
| aCTLA-4.9 | EIVLTQSPGTLSLSPGEGATLSCRASQSFSSNYLAWYQQKPGQAPRLLIYGASSRA<br>TGIPDRFSGSGSGTDFTLTISRLEPEDFAVYYCQQYGTSPFTFGPGTKVDIK | QVQLVESGGGVVQPGRSLRLSCAASGFTFSSFGMHWVRQAPGKGLEWVAVIWYDGRNKYYVDSVK<br>GRFTISRDNSNNTLYLQMNSLRAEDTAVYYCARGEFFGEFFDYWGQGTLVTVSSA |
| aCTLA-4.11 | EIVLTQSPGTLSLSPGERATLSCRASQSVSYLAWYQQKPGQAPRLLIYGASSRATG<br>IPDRFSGSGSGTDFTLTISRLEPEDFAVYYCQQYGSSPWTFGQGTKVEIK | QVQLVESGGGVVQPGRSLRLSCAASGFTFSSYGMHWVRQAPGKGLEWVAVIWYDGSNKFYADSVK<br>GRFTISSDNSKNTLYLQMNSLRAEDTAVYYCARGGHLGSFDYWGQGTLVTVSSA |
| aCTLA-4.12 | EIVLTQSPGTLSLSPGERATLSCRASQSVSSSYLAWYQQKPGQAPRLLIYGASSRA<br>TGIPDRFSGSGSGTDFTLTISRLEPEDFAVYYCQQYGTSPWTFGQGTKVEIK | QVQLVESGGGVVQPGRSLRLSCAASGFTFSSYGMHWVRQAPGKGLEWVAVIWYDGSNKYYADSVK<br>GRFTFSRDNSKNTLYLQMNSLRAEDTAVYYCARGGLMGAFDYWGQGTLVTVSSA |
| aCTLA-4.14 | EIVLTQSPGTLSLSPGERATLSCRASQSVSSSYLAWYQQKPGQAPRLLIYGASSRA<br>TGIPDRFSGSGSGTDFTLTISRLEPEDFAVYYCQQYGSSPFTFGPGTKVDIK | QVQLVESGGGVVQPGRSLRLSCAASGFTFSSYGMHWVRQAPGKGLEWVAVIWYDGRNKDYADSVK<br>GRITISRDNSKNTLYLQMNSLRAEDTAVYYCARGGLLGPYFDYWGQGTLVTVSSA |
| aCTLA-4.15 (GIGA-564) | EIVLTQSPGTLSLSPGERATLSCRASQSVSSSYLAWYQQKPGQAPRLLIYGASSRA<br>TGIPDRFSGSGSGTDFTLTISRLEPEDFAVYYCQQYGSSPWTSGQGTKVEIK | QVQLVESGGGVVQPGRSLRLSCAASGFTFSSYGMHWVRQAPGTGLEWVAVIWYEGRNKYYADPVK<br>GRFTISRDNSKNTLYLQMNSLRDDDTAVYYCARAGDLGAFDIWGQGTMVTVSSA |
| aCTLA-4.22 | EIVLTQSPGTLSLSPGERATLSCRASQSVSSSYLAWYQQKPGQAPRLLIYGASSRA<br>TGIPDRFSGSGSGTDFTLTISRLEPEDFAVYYCQQYGSSPFTFGPGTKVDIK | QVQLVESGGGVVQPGRSLRLSCVASGFTLSSYGMHWVRQAPGKGLEWVAVIWYDGSNKHYADSVK<br>GRFTISRDNSKNTLSLQMNSLRAEDTAVYYCARGGQLGPFDYWGQGTLVTVSSA |
| aCTLA-4.24 | DIQMTQSPSSLSASVGDRVTITCRASQSISSYLNWYQQKPGKAPKLLIYAASSLQS<br>GVPSRFSGSGSGTDFTLTISSLQPEDFATYYCQQSYSTPFTFGPGTKVDIK | QVQLVESGGGVVQPGRSLRLSCAASGFTFSSHGMHWVRQAPGKGLEWVAVIWYDGSNKHYADSVK<br>GRFTISRDNSKNTLYLQMNSLRAEDTAVYYCARGDILTGYYGYWGQGTLVTVSSA |
| aCTLA-4.26 | DIQMTQSPSSLSASVGDRVTITCRASQSISSYLNWYQQKPGKAPKLLIYAASSLQS<br>GVPSRFSGSGSGTDFTLTISSLQPEDFATYYCQQSYSTPFTFGPGTKVDIK | QVQLVESGGGVVQPGRSLRLSCAASGFTFSSYGMHWVRQAPGKGLEWVAVIWYDGSYKYYADSVK<br>GRFTISRDDSKNTLYLQMSSLRAEDTAVYYCARAPHYAILTGYYEDYWGQGTLVTVSSA |
| aCTLA-4.28 | EIVLTQSPGTLSLSPGDRATLSCRASQSGSSSYLAWYQQKPGQAPRLLIYGASSRA<br>TGIPDRFSGSGSGTDFTLTISRLEPEDFAVYYCQQYGTSPFTSGPGTKVDIK | QVQLVESGGGVVQPGRSLRLSCAASGFTFSSYGMHWVRQAPGKGLEWVAVIWYDGNNKYYADSVK<br>GRFTISRDNSKNTLYLQMYSLRAEDTAVYYCARGGILAAGIFDYWGQGTLVTVSSA |
| aCTLA-4.29 | EIVLTQSPGTLSLSPGDRATLSCRASQSGSSSYLAWYQQKPGQAPRLLIYGASSRA<br>TGIPDRFSGSGSGTDFTLTISRLEPEDFAVYYCQQYGTSPFTSGPGTKVDIK | QVQLVESGGGVVQPGRSLRISCAASGFTFSSYGIHWVRQAPGKGLQWVAVIWYDGRNKYYADSVK<br>GRFTISRDNSKNTLYLQMNSLRAEDTAVYYCSRSGSFGAFDIWGQGTMVTVSSA |

**Table S4. CTLA-4 antibody sequences.**

Amino acid sequences are provided of the IgK and IgG variable regions for the 14 characterized full-length CTLA-4 antibodies.
